# Attention enhances category representations across the brain with strengthened residual correlations to ventral temporal cortex

**DOI:** 10.1101/2021.12.15.472647

**Authors:** Arielle S. Keller, Akshay Jagadeesh, Lior Bugatus, Leanne M. Williams, Kalanit Grill-Spector

## Abstract

How does attention enhance neural representations of goal-relevant stimuli while suppressing representations of ignored stimuli across regions of the brain? While prior studies have shown that attention enhances visual responses, we lack a cohesive understanding of how selective attention modulates visual representations across the brain. Here, we used functional magnetic resonance imaging (fMRI) while participants performed a selective attention task on superimposed stimuli from multiple categories and used a data-driven approach to test how attention affects both decodability of category information and residual correlations (after regressing out stimulus-driven variance) with category-selective regions of ventral temporal cortex (VTC). Our data reveal three main findings. First, when two objects are simultaneously viewed, the category of the attended object can be decoded more readily than the category of the ignored object, with the greatest attentional enhancements observed in occipital and temporal lobes. Second, after accounting for the response to the stimulus, the correlation in the residual brain activity between a cortical region and a category-selective region of VTC was elevated when that region’s preferred category was attended vs. ignored, and more so in the right occipital, parietal, and frontal cortices. Third, we found that the stronger the residual correlations between a given region of cortex and VTC, the better visual category information could be decoded from that region. These findings suggest that heightened residual correlations by selective attention may reflect the sharing of information between sensory regions and higher-order cortical regions to provide attentional enhancement of goal-relevant information.

## I. Introduction

In the natural world, the human visual system is constantly inundated by many competing stimuli, some of which are relevant for behavioral goals and others that are irrelevant. To sort through this abundant visual input, human observers selectively focus their attention on important or goal-relevant stimuli, while ignoring irrelevant distractions, a process known as selective attention. However, it remains a mystery how the neural representations of attended and ignored items change to facilitate selective processing of relevant stimuli. Solving this ongoing puzzle requires understanding how visual inputs are represented across the brain as well as understanding how large-scale networks coordinate the allocation of attention to relevant representations.

It is well-documented that attention improves performance on a wide variety of tasks such as sensory discrimination (Lee et al., 1997) and target detection (Posner, 1980). The question of *how* attention leads to improved behavioral performance has been a subject of prior research in both humans and animal models for many years revealing that: attention increases neural firing rates (Motter, 1993), tunes visual cortical responses (Desimone and Duncan, 1995; Kastner et al., 1999, 1998), and is associated with coordinated activity in a large-scale fronto-parietal network (FPN; Corbetta and Shulman, 2002; Kastner et al., 1999; Nobre et al., 1997) that is related to goal-directed attention abilities (Fellrath et al., 2016; Prado et al., 2011). Although such studies have laid the groundwork for understanding the neural correlates of attention, we lack a clear understanding of how attending to or ignoring sensory stimuli modulates their neural representations across the brain.

Prior research on visual attention in humans has either examined how attention modulates the amplitude of cortical responses (Kay and Yeatman, 2017; Wojciulik and Kanwisher, 1999) and cortical representations of visual stimuli (Baldauf and Desimone, 2014; Bugatus et al., 2017; Córdova et al., 2016; Çukur et al., 2013; Peelen et al., 2009) or has examined how attention affects the interaction between ongoing activity in various brain regions (Al-Aidroos et al., 2012; Chadick and Gazzaley, 2011; Norman-Haignere et al., 2012) during an attentionally demanding task. The former research examined the effect of attention on bottom-up stimulus-evoked responses focusing mainly on high-level visual cortex in lateral occipital-temporal cortex (LOTC) and ventral temporal cortex (VTC) where category-selective regions reside (Kanwisher, 2010) and distributed visual category representations are salient across large cortical expanses (Cox and Savoy, 2003; Haxby et al., 2001; Kriegeskorte et al., 2008; Weiner and Grill-Spector, 2013). These studies revealed that visual attention to items of certain categories enhances responses in category-selective regions of the attended category (Kay and Yeatman, 2017; Moore et al., 2013; Wojciulik and Kanwisher, 1999) as well as the distributed representations (Çukur et al., 2013; Peelen et al., 2009). Nonetheless, other studies revealed that distributed responses in LOTC and VTC represent category information for both attended and unattended items (Bracci and Op de Beeck, 2016; Bugatus et al., 2017).

Other research examined how attention affects the interaction between brain areas by measuring the correlations between the residual activity of pairs of brain areas after accounting for the stimulus driven component, as the residual ongoing activity is thought to capture more of the top-down activity that one might continuously maintain while performing a task and is not locked to the stimulus. This approach, which we refer to as “residual correlations,” measures the correlation between residual activities across brain regions and has been referred to elsewhere as “background connectivity” (Al-Aidroos et al., 2012) or “task-residual functional connectivity” (Tran et al., 2018). Prior research has revealed that attention to visual items increases the strength of residual correlations between cortical regions of the frontal parietal network (FPN) and visual cortex (Chadick and Gazzaley, 2011; Griffis et al., 2015) as well as between category-selective regions of VTC (Norman-Haignere et al., 2012).

However, two main gaps in knowledge remain. First, it is unclear whether selective attention influences representations of visual object categories across the entire brain as most prior studies have focused on a handful of theoretically important, predefined cortical regions (but see Çukur et al., 2013). Second, it remains unknown whether neural representations of attended or ignored information vary with the strength of residual correlations between cortical areas and sensory regions processing the attended or ignored information.

To address these gaps in knowledge we examined the effect of attention on both visual category representations across the entire cortex and residual correlations between each cortical region in the brain and category-selective regions in VTC (Kanwisher, 2010; Peelen and Downing, 2005), when their preferred category is selectively attended or ignored. To do so, we leveraged a selective attention task we previously developed (Bugatus et al., 2017) while participants underwent fMRI scanning. In this task (Fig 1), participants were asked to view superimposed images of two categories, attend to items of one category and indicate when items of the attended category were inverted.

**Figure 1:**
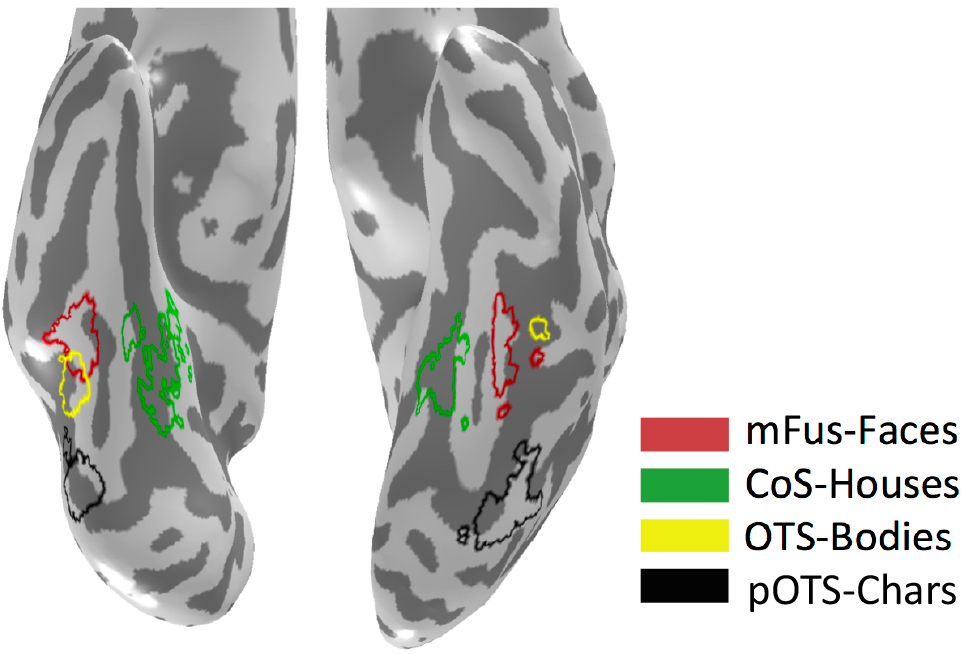
Example VTC category selective regions. VTC category-selective regions in an example participant.

This experimental paradigm has three key advantages. First, this selective attention task utilized stimuli from five visual object categories that can be readily decoded from their distributed responses across LOTC and VTC (Bracci et al., 2017; Bracci and Op de Beeck, 2016; Bugatus et al., 2017; Cox and Savoy, 2003; Grill-Spector and Weiner, 2014; Haxby et al., 2001; Kriegeskorte et al., 2008; Proklova et al., 2016), suggesting that we can use these stimuli to measure the effect of attention on bottom-up category representations for attended and ignored stimuli. Second, we use stimuli that are associated with category-selective regions in VTC (Kanwisher, 2010), in which activity is critical for recognition of these categories (Gaillard et al., 2006; Jonas et al., 2014; Parvizi et al., 2012). This allowed us to examine the effect of attention on top-down brain activity by measuring residual correlations between each brain region and category-selective region when their preferred category was attended or ignored. Third, images of items from these attended and ignored object categories were overlaid in a single spatial location, and subjects were asked to conduct a perceptual task, which allowed us to probe the effects of attending and ignoring during visual competition without shifting the spatial focus of attention and without necessitating other cognitive processes such as working memory.

Additionally, we utilized a unique analytical approach, combining two quantitative techniques to answer our three main questions of interest. First, we aimed to test how selective attention modulates the representation of attended vs. ignored visual categories by testing its influence on the decodability of visual category information when the same items are attended or ignored. We used a data-driven whole-brain approach to determine where in the brain information about attended and ignored visual object categories are decodable. To do so, we examined the classification accuracy of category information in each of the 180 brain areas of the Glasser atlas (Glasser et al., 2016), which is the most recent brain parcellation based on both functional and structural metrics. While it could be the case that information about attended and ignored object categories are equivalently decodable in all areas of cortex, or that attentional enhancement is present in all areas of cortex, based on prior findings summarized above, we hypothesized that both category information and the amount of attentional enhancement would vary across cortical regions.

Second, we aimed to test in a data-driven, whole-brain manner which cortical regions have strong residual correlations with category-selective regions of VTC when the latter regions’ preferred categories are either attended or ignored. Using functionally-defined category-selective regions of VTC representing either attended or ignored object categories provided us with an anchor to then investigate residual correlations between these regions and other parts of the brain. We hypothesized that the strongest residual correlations would be observed between regions of cortex and a particular category-selective region of VTC (e.g., a face-selective region in the fusiform gyrus) when participants are cued to attend that region’s preferred category (e.g., faces). However, an alternative hypothesis, consistent with literature showing that both attending and ignoring require top-down cognitive control (Martinez-Trujillo and Treue, 2004; Scolari et al., 2012), is that residual correlations with category-selective regions would be equally strong regardless of whether those preferred categories are attended or ignored. By utilizing both actively-attended and actively-ignored stimuli presented simultaneously in the same spatial location, we directly pit these competing hypotheses against one another.

Third, importantly, we investigated the relationship between the effects of attention on stimulus representations and residual correlations. That is, we sought to determine (i) whether there is a correlation between a region’s category representations and its residual correlations with VTC category-selective regions, and (ii) whether this relationship is modulated by attention. First, we hypothesized that if residual correlations reflect the sharing of information about attended visual object categories between VTC and other cortical regions, then it should follow that category classification accuracy would be higher in cortical regions with stronger residual correlations with VTC category-selective regions. Second, we hypothesized that selective attention would modulate this relationship, revealing a stronger correlation between residual correlations and classification accuracy when images from these object categories are attended compared to when they are ignored. In particular, we hypothesized that this attentional enhancement of classification accuracy and residual correlations would be prominent in regions of the FPN, given this network’s established role in goal-directed attention. Together this combined novel approach allows us to systematically examine the impact of top-down selective attention on object category representations and their relationship across the brain.

## 2. Materials and Methods

The fMRI data presented here were previously published in (Bugatus et al., 2017) and (Keller et al., 2021). Here we will briefly describe the subjects and acquisition as detailed information can be found in the original manuscript. Additionally, we describe in detail new analyses and methodological approaches that are unique to the present paper and have not been published or done elsewhere. **Data/Code Availability Statement:** Code for our key analyses is available at https://github.com/akjags/att_class_resid. Data may be made available by requests from the authors.

### 2.1 Subjects

Subjects recruited from Stanford University participated in one of two studies. The first study included twelve participants (5 female, ages 23-44) whose data was previously published in (Bugatus et al., 2017). These participants underwent fMRI scanning including three runs of the Selective Attention task and three runs of the Oddball task. The second study included 20 additional participants (10 female, ages 18-37) from (Keller et al., 2021) who participated in one run of the Selective Attention task and one run of the Oddball task as well as other tasks not relevant to this study. Because of time constraints, they participated in fewer runs of the Selective Attention and Oddball tasks. Seven subjects were excluded because of excessive head movement (>2 voxels either within-scan or between-scans) during one or more tasks and 4 subjects were excluded because we could not localize the majority of VTC functional ROIs of sufficient size. Thus, a total sample of twenty-one subjects (8 female, ages 21-44) were included in our analyses. All subjects had normal or corrected-to-normal vision. **Ethics Statement:** All procedures were approved by the Stanford Internal Review Board on Human Subjects Research. Participants gave written informed consent before participating in this study.

### 2.2 Data Acquisition and Preprocessing

Subjects were scanned using a General Electric Sigma MR750 3T scanner located in the Center for Cognitive and Neurobiological Imaging (CNI) at Stanford University using a custom-built 32-channel head coil. Using an EPI sequence with a multiplexing (multiband) factor of 3, we acquired 48 slices at 2.4 mm isotropic resolution, FOV = 192 mm, TE = 30 ms, TR = 1 s, and flip angle = 62°. The slice prescription covered the entire brain, except for the very superior portion of the cortex, roughly corresponding to superior motor and somatosensory cortices. Additionally, T1-weighted anatomicals of the same prescription were acquired, which were used to align the fMRI data to the whole brain anatomical images. Finally, whole-brain anatomical images of each subject’s brain were acquired using a T1-weighted SPGR sequence with a resolution of 1×1×1mm, FOV = 240mm, flip angle = 12°. This volume anatomy was used to create a cortical surface reconstruction of each subject’s brain using FreeSurfer 6.0 (https://surfer.nmr.mgh.harvard.edu/).

Functional data from each run was aligned to the individual’s own brain anatomy. Motion correction was performed both within and across functional runs using mrVista (https://github.com/vistalab/vistasoft) motion correction algorithms. Runs in which participants’ head motion was greater than 2 voxels were discarded. No slice-timing correction or global signal regression was performed. All data were analyzed within individual participants’ native brain anatomy space without spatial smoothing.

### 2.3 Selective Attention Task

During fMRI, subjects viewed grayscale images from various object categories: faces, houses, cars, bodies, and pseudo-words while fixating at the center of the screen. Examples of the stimuli and task are in (Bugatus et al., 2017). In each 8-second block, subjects were presented with a series of eight grayscale images each containing overlaid exemplars from two object categories (e.g., faces and houses). Before each block, a cue indicating the name of the category to be attended appeared for 1 second. Participants were instructed to attend to items of that category (e.g., faces), and to respond with a button press when an item of the attended category was presented upside-down. 0, 1, or 2 images of either the attended or ignored category were presented upside down at random in each block. This paradigm also necessitates active feature-based ignoring as subjects were instructed to withhold responses to upside-down items of the Ignored category that occurred in the same frequency as the attended one. Subjects performed between one and three runs of this task with different images. Image order and presentation was randomized across runs. Each run contained 40 blocks. Across blocks, all possible pairings of the 5 categories and attended/ignored conditions occurred.

### 2.4 Oddball Task

The same participants also performed in a fMRI experiment in which they performed an oddball detection task. In this task, participants viewed grayscale images from the same categories: faces, houses, cars, bodies, and pseudo-words. Fixating at the center of the screen, subjects are presented with a series of 8 images in each block and were asked to respond when a phase-scrambled image without an object appeared. Either 0, 1 or 2 phase-scrambled images were presented in each 8 image block. Since each item was presented individually, the neural responses measured during this task were used as the training data for the classification analysis.

### 2.5 Definition of regions of interest (ROIs)

We used two types of regions of interest (ROIs) to analyze the data: Functional ROIs (fROIs) to define category-selective regions of VTC at the individual subject level, and Glasser Atlas ROIs (Glasser et al., 2016) that tile the entire cortex.

#### 2.5.1 Functional Regions of Interest (fROIs)

To independently identify category-selective regions of VTC, we used an independent localizer experiment in which subjects viewed in blocks images from 5 domains (faces, bodies, places, characters and objects). The localizer used 3 runs, similar to the (Stigliani et al., 2015) (experiment is available here: https://github.com/VPNL/fLoc). 8 subjects participated in a localizer with an oddball task and 2 categories per domain (as in Stigliani et al., 2015) and 24 subjects participated in an experiment with images from the same 5 domains (1 category per domain) and a 2-back task (as in Bugatus et al., 2017). Prior experiments from our lab show that both tasks are able to localize VTC category-selective regions effectively (Bugatus et al., 2017; Weiner and Grill-Spector, 2010). Therefore, as a logistical convenience, the *n*=8 participants who had already participated in an Oddball task localizer experiment (and whose fROIs had already been successfully identified) were not asked to complete another localizer experiment. This did not impact our ability to identify fROIs in all of the participants.

We analyzed the localizer data using a general linear model (GLM): (1) Using the GLM we estimated block-averaged response amplitudes (betas) to each category in each voxel. (2) Then, we generated several contrast maps comparing responses to images of one domain vs. all other domains (units of t-values). (3) Category-selective ROIs were defined as voxels in ventral temporal cortex (VTC) having significantly stronger responses to images of that domain compared to all others, with a threshold of t≥3 (as t represents effect size), voxel level, uncorrected. All fROIs were a minimum of 5mm^3^ and maximum of 177mm^3^ each, with an average volume of 43.4 ± 6.67mm^3^.

Here we analyzed data from one VTC category-selective region from each domain (faces, body, place, word) to test if effects vary across domains. As there are multiple face and word-selective ROIs in VTC, we selected one category-selective region within VTC for each object category of interest (faces, houses, bodies, words). This allowed us to compute the average residual correlation across conditions evenly, using one region per category for all categories. We chose mFus-Faces because it is anatomically proximal to OTS-bodies and CoS-places (e.g., Stigliani 2015), thought to be in the same level of the processing hierarchy (Weiner et al., 2017), and we were able to identify these regions in the majority of our participants. As mOTS-chars is left lateralized (Gomez et al., 2018; Stigliani et al., 2015) (only ~20% of participants have bilateral mOTS) but pOTS-chars is found bilaterally, we chose to use pOTS-chars rather than mOTS-chars as the other category-selective fROIs are bilateral. Supplementary Table 1 provides the details as to which fROI was identified in each participant and hemisphere and Figure 1 shows example fROIs.

##### mFus-faces

was defined as a cluster of face-selective voxels in the lateral fusiform gyrus near or overlapping the anterior tip of the mid fusiform sulcus (MFS), as in (Weiner et al., 2017). We identified mFus-faces in 19/21 subjects in the right hemisphere and 14/21 subjects in the left hemisphere.

##### CoS-places

was defined as a cluster of place-selective voxels in the collateral sulcus (CoS) near/overlapping the junction between the CoS and the anterior lingual sulcus (ALS) as in (Weiner et al., 2017). We identified CoS-places in 21/21 subjects in the right hemisphere and 21/21 subjects in the left hemisphere.

##### OTS-bodies

was defined as a cluster of body-selective voxels in the occipital temporal sulcus (OTS), typically between pFus- and mFus-faces (Weiner et al., 2017; Weiner and Grill-Spector, 2010). We identified OTS-bodies in 19/21 subjects in the right hemisphere and 10/21 subjects in the left hemisphere.

##### pOTS-chars

was defined as a cluster of character-selective voxels in the OTS. We identified pOTS-chars in 13/21 subjects in the right hemisphere and 17/21 subjects in the left hemisphere.

#### 2.5.2 Glasser Atlas ROIs

To independently define ROIs tiling the entire cortex, we used the Glasser Atlas (Multi-Modal Human Connectome Project’s Atlas (HCP-MMP1.0)) (Glasser et al., 2016)). We chose this brain parcellation because (i) it is the most up to date parcellation of the entire brain and (ii) it is based both on functional and connectivity properties of cortical regions which makes it unbiased and appealing for the questions of interest in the present study. We used the ROIs defined in the FreeSurfer 6.0 (https://surfer.nmr.mgh.harvard.edu/) average brain (labels in FreeSurfer) and using cortex-based alignment in FreeSurfer we transformed these labels into each participant’s native cortical surface. Then we imported each ROI from FreeSurfer into mrVista for subsequent functional analysis. No additional voxel selection or ROI restriction was performed. Because ROI size is non-uniform in the Glasser atlas varies, we tested whether our key measurements (classification accuracy, Methods Section 2.6; and residual correlations, Methods Section 2.7) are correlated with ROI size, which we computed as the number of voxels contained in the ROI on the freesurfer average (fsaverage) inflated surface. Results shown in Supplementary Table 2 reveal no correlation between classification accuracy or residual correlations and Glasser ROI size.

For analyses in which ROIs were analyzed separately by lobe, we assigned each Glasser Atlas ROI to its most proximal lobe: occipital, temporal, parietal, or frontal. Lobe assignments for each Glasser Atlas ROI are depicted in Supplementary Figure 1. The following Glasser atlas ROIs which overlap the average VTC functional ROIs described above (Rosenke et al., 2021; Weiner et al., 2017) were excluded from all statistical analyses: FFC, PH, PHA1, PHA2, PHA3, TE2p, VMV1, VMV2, VMV3. These ROIs are depicted in Supplementary Figure 2. Based on our hypothesis that regions of the fronto-parietal network (FPN) involved in goal-directed attention might show particularly notable enhancement of classification accuracy (Methods Section 2.6) and residual correlations (Methods Section 2.7) with attention, we made note of the following Glasser Atlas ROIs that overlap the FPN (Osher et al., 2019): 6a, 6ma, 6d, 6v, 6r, i6-8, 8Av, 8C, 55b, FEF, PEF, IFJp, IFJa, IFSp, V7, IP0, IP1, IPS1, IP2, AIP, LIPd, LIPv, VIP, MIP, 7PL, 7PC, 7AL, and PFt. These ROIs are also depicted in Supplementary Figure 2.

### 2.6 Category Classification from Cortical Responses

To assess the degree to which category information is represented across cortex, we employed a multi-class support vector machine classifier (SVM, Boser et al., 1992), by constructing multiple one-versus-all SVM classifiers with a linear kernel to predict the object category that was attended or ignored from multivoxel patterns of activity. The SVM was trained and tested on distributed responses across voxels of each of the Glasser Atlas ROIs. Other hyperparameter values include box constraint of 1, kernel scale of 1, alpha initialization of 0.5, and a Sequential Minimal Optimization (SMO) solver.

#### Classification Procedure

Classification was done independently in each ROI from the distributed responses of that ROI. The training and testing sets were from independent experiments: the training set consisted of data from the Oddball task and the test sets were the Attended or Ignored conditions during the Selective Attention task. We assessed the performance of the classifier that had been fit to the Oddball task in predicting the attended or ignored category in the Selective Attention task. Prior research suggests that category-specific information from raw response amplitudes (betas) can be cloaked by the mean response across all categories (Margalit et al., 2020; Sayres and Grill-Spector, 2008). Thus, in the classification analyses, we used t-contrasts to minimize the shared variance in the cortical responses to different categories (e.g., distance from the coil) and to examine the relative contribution of a category relative to other categories while taking into account the residual error of the GLM.

#### Training

The training set for each ROI consisted of the distributed responses from the Oddball tasks across the ROI, which were based on t-values. In brief, we first fit data from the Oddball task in each voxel in the brain using a GLM by convolving the design matrix of the Oddball experiment with the hemodynamic response function (hRF) implemented in SPM8 (https://www.fil.ion.ucl.ac.uk/spm/), to estimate response amplitudes (betas) for each of the five categories. From the GLM, we computed in each voxel a t-contrast for each category against all other categories. We extracted the distributed pattern of t-values across voxels in a given ROI and used this to train the classifier.

#### Testing

The testing set consisted of data in the same ROI from the Selective Attention task. Like the Oddball task, we fit a GLM in each voxel to the data of the Selective Attention task, and estimated response amplitudes (betas) to each of the 20 conditions of the task (attend/ignored x 10 pairings of 5 categories). From this we estimated distributed responses across the ROI for the attend/ignore conditions for each category. <u>Attended</u>: After fitting the GLM to the data of the Selective Attention task, we computed the t-contrast in each voxel, contrasting all the conditions in which a given category was attended against all the conditions in which that category was not present. This results in 5 contrasts. We then extracted the distributed pattern of t-values across voxels in a given ROI to evaluate the classifier. <u>Ignored</u>: Same as attended, but for all conditions in which the category was present and ignored vs. all conditions in which the category was not present. We performed these processing steps separately for each of the 3 runs, giving us a total of 15 observations for each of our test sets.

Using the data from the Oddball task as SVM training data, we tested the performance of the SVM classifier separately on the attended and ignored conditions. That is, in the attended condition the classifier predicts what category the subject is attending to and in the ignored condition the classifier predicts which category the subject is ignoring.

In order to perform multiclass classification with a linear SVM, we employed the Matlab (mathworks.com) functions fitcsvm and fitcecoc to train a binary classifier for each category (1 vs. all others) and generate a prediction based on which classifier had the greatest separation from its category boundary to determine the category out of the 5 possible categories. We computed the noise ceiling performance by randomly permuting the category labels and using the trained multi-class SVM classifier to predict the randomly permuted labels 1000 times. This gave us a distribution of prediction accuracies, from which we could estimate the threshold required to exclude 95% of these predictions. This was computed individually within each subject and each ROI separately, and then averaged together to yield a global threshold that could be compared across all ROIs. However, we also verified that computing the confidence intervals separately and thresholding each ROI using a ROI-specific threshold did not change the number of ROIs which exceeded chance level classification accuracy in the brain maps. We thus used an accuracy threshold of 0.2002 ± 0.0234 to decide which ROIs exceeded chance level classification.

#### Classification control

To test whether classification results are specific to this dependent variable (t-values), we replicated these classification analyses using raw responses (betas) for both the training and testing data. Results in Supplementary Figure 3.

### 2.7 Residual Correlation Analysis

To uncover functional relationships associated with top-down signals between cortical regions and category-selective regions under attended and ignored conditions of the Selective Attention task, we performed a residual correlation analysis (Supplementary Figure 4). First, we sought to isolate the top-down component of the BOLD signal that is independent from the bottom-up, stimulus driven component. Thus, we separated each voxel’s time course to 2 components: (i) the stimulus-evoked component, and (ii) the residual activity. To estimate the stimulus-evoked activity, we fit a general linear model (GLM) to the time course by convolving the experimental design matrix with the hemodynamic response function to generate predictors of the contribution of each condition to the BOLD response. Fitting each voxel’s time course data, we estimated betas for each predictor separately for each run. Then we extracted the residual activation in each voxel by subtracting the predicted time course calculated from the GLM from the measured voxel time course.

After these whole time course residuals were computed, we extracted the residual for each trial type: that is, each of four stimulus categories which are associated with a VTC category-selective ROI (faces, houses, bodies, and words) separately for when it was presented and attended (e.g., trials in which faces were present and participants were cued to attend to faces) and when it was presented and ignored (e.g., trials in which faces were present but a different category was attended to) and these trials (40 volumes from each condition per run) were concatenated. Then, we calculated mean residuals across voxels of each of the VTC category-selective fROIs (mFus-Faces, CoS-Houses, OTS-Bodies, OTS-Words) as well as each Glasser Atlas ROI to determine the mean residual of each ROI. Since averaging across voxels removes independent noise among voxels, this residual reflects a component of the brain signal that is not explained by the stimulus, for example, top-down attention is not modeled in the GLM. To determine the correlation in the time-series of residuals between ROIs, we then calculated the pairwise correlations between the average residual of each category selective ROI and each Glasser Atlas ROI. Correlations were calculated separately for trials in which the preferred stimulus category for each ROI (e.g., faces for mFus-Faces) was attended and when it was ignored. We refer to these resulting correlations as “residual correlations,” as they refer to correlation between the residual signals of pairs of ROIs. Others have referred to these correlations as “background connectivity” (Al-Aidroos et al., 2012).

To test whether the GLM captured the stimulus-evoked activity, we performed a control analysis, in which we computed correlations in residual activity between two trials of the same condition (e.g., face attended, body ignored condition). This control was done within each region of the Glasser Atlas in each hemisphere and subject (for this analysis, we utilized the 12 subjects who underwent three total runs of the experiment to maximize the number of trials of each condition type). We reasoned that if after removing the stimulus-evoked activity estimated by the GLM, there still remained some stimulus-related activity that was not modeled by the GLM (e.g., offset response), then the within-region residual correlations among trials of the same condition would be significantly positive. However, contrary to this prediction, results of this analysis show that the distribution of these within-ROI residual correlations to the same condition is not significantly greater than zero (mean ± std: −0.010±0.051; *t*(11)=-0.712, *p*=0.492, two-sided). As a second control, we conducted the same procedure on the original time series prior to removing the activity modeled by the GLM. Results show that doing the same analysis on the original data (prior to removing the task-based activation from the GLM) the mean within-region correlation to different trials of the same condition was positive on average (mean±std: 0.008±0.044) and significantly greater than the distribution generated using the residual correlations (*t*(11)=3.069, *p*=0.011, two-sided; Supplementary Figure 5). These analyses provide strong evidence that the GLM procedure effectively captures the stimulus-evoked activity. Residual correlations were computed using Matlab R2014a (mathworks.com), mrVista (https://github.com/vistalab/vistasoft), and SPM8 (https://www.fil.ion.ucl.ac.uk/spm/software/spm8/).

#### Chance-Level Residual Calculations

To determine chance level residual correlation for each analysis, we used permutation testing. We hypothesized that if the residuals reflect top-down signals then they should be subject-unique. However, if there remained task-based activity that was not removed by the GLM, it would have remained across subjects and thus would have elevated the chance level residual correlations between subjects. Thus, for each Glasser Atlas ROI, we calculated correlations between the residuals in that ROI in one subject with the residuals of a VTC category-selective ROIs for a randomly-selected condition from a different subject. This process was repeated over 1000 iterations for each Glasser Atlas ROI, each time randomly choosing two independent subjects, a VTC category-selective fROI, and a condition by sampling randomly with replacement. The mean correlation across these 1000 iterations provides an estimate of the chance residual correlation which was 0.01±0.03. To further ensure that this chance-level was a reasonable estimate, we also computed a second chance-level aimed at breaking the temporal correlations between residuals: we randomly shuffled the time series of residuals in each region and subject before computing pairwise correlations between residuals of different ROIs. This random shuffling procedure was repeated 1000 times for each region of the Glasser Atlas, in each hemisphere, and subject. Then we calculated the average chance level for each subject by averaging over regions and hemispheres and confirmed that no subjects’ shuffled chance-level were outliers (greater than two standard deviations of the mean). We then calculated the average chance-level across all subjects (−0.00002) as well as the 95th percentile (0.017). Given that this second chance-level was less stringent than the above chance-level, we opted to use the stricter chance-level as a reference point in our figures.

### 2.8 Statistical Analyses

To assess the significance of classification accuracy across Glasser ROIs and residual correlations of Glasser ROIs with VTC ROIs, we utilized separate three-way repeated measures analyses of variance (ANOVAs) for each metric with factors for lobe (frontal, parietal, temporal and occipital), hemisphere (left or right), and condition (attended or ignored) totaling 4×2×2 factors. To determine whether the effects of selective attention on residual correlations and classification accuracy varied systematically by their magnitude, we performed a regression analysis relating the magnitude of the metric (classification accuracy/residual correlations) separately for the attended and ignored conditions across ROIs in each lobe. The slope of the regression can be thought of as an attentional scaling factor, which represents the extent to which attention scales up or scales down each metric; Tables 1 and 2, respectively, provide details of linear regression results, with with *t* and *p* values indicating whether the coefficients of the linear regression are significantly greater than zero (two-sided test). We applied Bonferroni correction for multiple comparisons to account for the eight linear models applied for each metric (classification accuracy and residual correlations). All statistical analyses were conducted in MATLAB 2014a (mathworks.com) and R (Version 3.5.0) using RStudio (Version 1.1.383).

**Table 1.** Linear regression results relating the average classification accuracy in each lobe and hemisphere during attended and ignored conditions. N represents the number of ROIs per lobe. Bold *p*-values represent significant attentional scaling after Bonferroni correction for 8 comparisons.

| Lobe | Hemisphere | N | Attentional Scaling | Std. Error | t | p |
| --- | --- | --- | --- | --- | --- | --- |
| Occipital | Left | 18 | 1.47 | 0.21 | 6.94 | **3.35 x 10<sup>-6</sup>** |
|  | Right | 18 | 2.02 | 0.23 | 8.82 | **1.52 x 10<sup>-7</sup>** |
| Temporal | Left | 33 | 1.92 | 0.21 | 9.30 | **1.77 x 10<sup>-10</sup>** |
|  | Right | 33 | 1.60 | 0.19 | 8.26 | **2.52 x 10<sup>-9</sup>** |
| Parietal | Left | 47 | 1.24 | 0.25 | 5.01 | **8.89 x 10<sup>-6</sup>** |
|  | Right | 47 | 1.14 | 0.20 | 5.64 | **1.06 x 10<sup>-6</sup>** |
| Frontal | Left | 73 | 0.17 | 0.13 | 1.34 | 0.184 |
|  | Right | 73 | 0.27 | 0.13 | 2.17 | 0.034 |

**Table 2.** Linear regression results relating the mean residual correlations in the attended and ignored conditions in each lobe and hemisphere. N represents the number of ROIs per lobe. Bold *p*-values represent significant attentional scaling after Bonferroni correction (8 comparisons).

| Lobe | Hemisphere | N | Attentional Scaling | Std. Error | t | p |
| --- | --- | --- | --- | --- | --- | --- |
| Occipital | Left | 18 | 1.18 | 0.10 | 11.56 | **3.52 x 10<sup>-9</sup>** |
|  | Right | 18 | 0.96 | 0.10 | 9.88 | **3.26 x 10<sup>-8</sup>** |
| Temporal | Left | 33 | 0.98 | 0.05 | 18.91 | **< 2.00 x 10<sup>-16</sup>** |
|  | Right | 33 | 0.93 | 0.06 | 16.35 | **< 2.00 x 10<sup>-16</sup>** |
| Parietal | Left | 47 | 1.25 | 0.08 | 15.66 | **< 2.00 x 10<sup>-16</sup>** |
|  | Right | 47 | 0.99 | 0.05 | 18.44 | **< 2.00 x 10<sup>-16</sup>** |
| Frontal | Left | 73 | 0.91 | 0.06 | 16.12 | **< 2.00 x 10<sup>-16</sup>** |
|  | Right | 73 | 0.91 | 0.04 | 22.92 | **< 2.00 x 10<sup>-16</sup>** |

Having computed the classification accuracy as well as the residual correlation for each ROI of the Glasser Atlas, we examined whether these measures were related. To quantify the relationship, we computed linear regressions between the vector of classification accuracies and the vector of residual correlations with VTC ROIs across the ROIs in each hemisphere of the Glasser Atlas. The linear regressions were computed between the mean subject classification performance for that ROI and the mean subject residual correlation with the VTC ROIs. The regression analysis was done separately in each hemisphere, lobe, and attentional condition (attended, ignored) and these 16 regressions were Bonferroni corrected to account for multiple comparisons. To test whether these linear relationships varied significantly by attention condition, we used stepwise linear regression to predict classification accuracy in each hemisphere and condition using residual correlations (step 1), attention condition (step 2), and the interaction between residual correlations and attention condition (step 3). Improvements in model performance at each step were calculated with a Chi-Squared test using the “anova” procedure in R.

## 3 Results

### 3.1 Does attention affect the decodability of category information in the human brain?

To examine the effect of attention on category representations across cortex, we compared a classifier’s ability to decode category information from brain responses during the Selective Attention task when a category was attended to when it was ignored. We decoded this information independently in each ROI of the Glasser Atlas (Glasser et al., 2016) and compared this to chance level classification. To that end, we used a support vector machine (SVM) classifier with a linear kernel to predict from each region’s distributed responses either the attended object category or the simultaneously presented Ignored object category during each trial of the Selective Attention task (Methods, section 2.6). Importantly, the classifier was trained on distributed responses from a separate Oddball task, in which the subjects viewed single stimuli rather than overlaid stimuli, with no attentional cues.

Qualitatively, we observed that the category of the attended stimulus can be decoded with accuracy above chance in many regions across the cortex, with the highest decoding accuracy found in visual cortex, including early visual cortex, lateral occipital cortex (LOC), and ventral temporal cortex (VTC) (Fig 2a). Glasser ROIs with notable classification accuracy (greater than 0.5 compared to 0.2 chance level) during the attended condition were: bilateral V2, V3, V3CD, V4, V4t, V8, LO1, LO2, LO3, PIT, and left hemisphere V1 and V3B in the occipital lobe, as well as bilateral VVC and right hemisphere FST in the temporal lobe (Fig 2a). We observed that several of the Glasser ROIs overlapping the fronto-parietal network (Supplementary Figure 2) were in the top 15% of ROIs with the highest enhancement of classification accuracy with attention: 7PL, IP0, IPS1, V7, and VIP. Interestingly, we also found that category information was decodable above chance in regions not typically associated with visual representation, including primary somatosensory cortex and primary motor cortex (BA2 and BA4). Furthermore, we found that the category of the ignored stimulus could also be decoded with accuracy above chance in several regions, particularly in Glasser ROIs spanning the visual cortex in the occipital and temporal cortices (Fig 2b).

**Figure 2:**
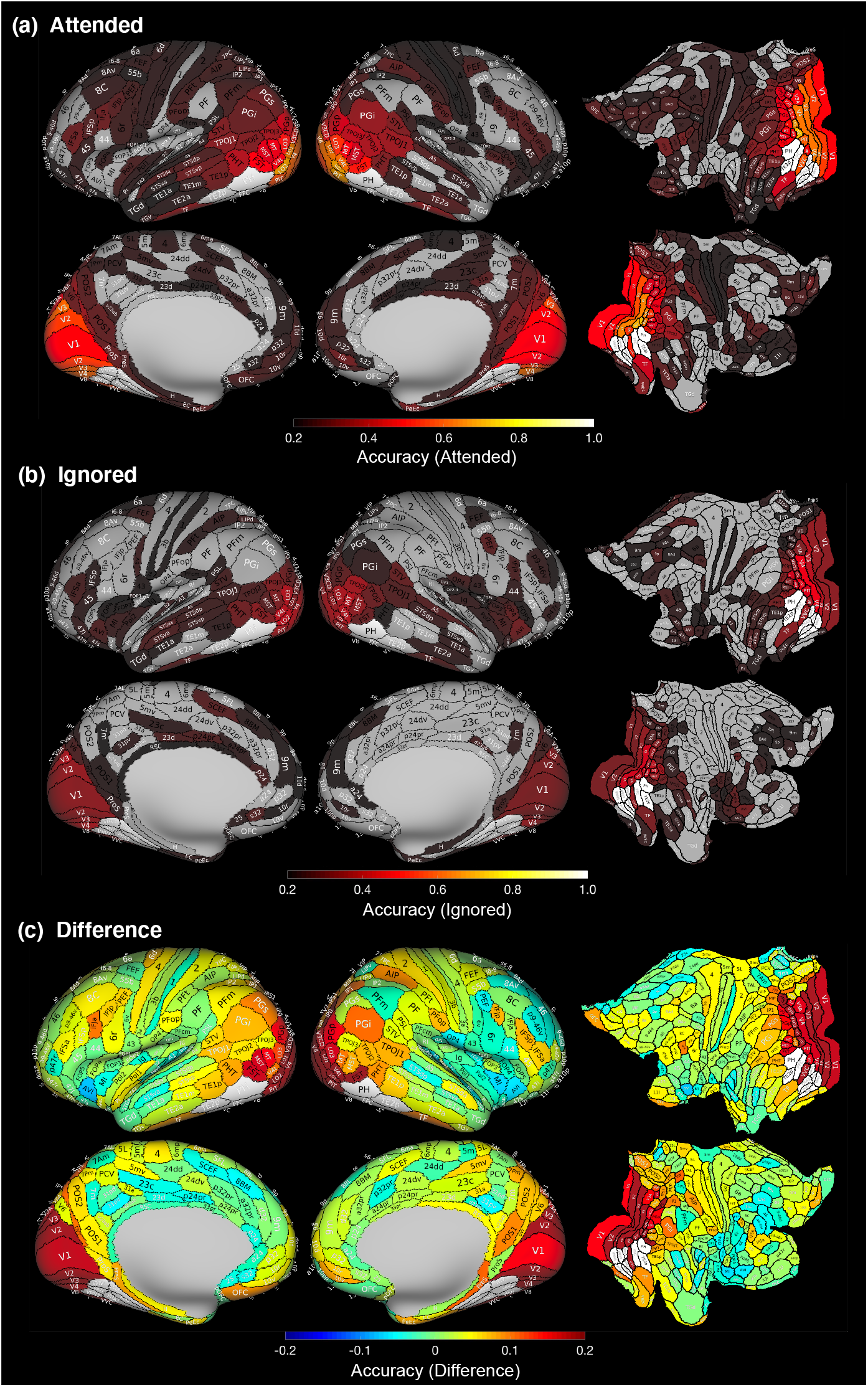
Classification of category information is higher for attended than ignored stimuli. Mean classification accuracy (proportion of object categories correctly classified) across 21 subjects and 5 object categories for (a) attended and (b) ignored categories. Maps are shown for the inflated lateral (top) and medial (bottom) cortical surfaces as well as flattened views (right) and are thresholded at chance level. That is, gray ROIs represent those with classification accuracy below chance level (0.2). (c) Mean difference in classification accuracy between attended and ignored conditions. Statistical significance values of the difference for each Glasser ROI are shown in Supplementary Figure 6a.

To better visualize where in the cortex category classification accuracy is enhanced by selective attention, we subtracted in each of the Glasser ROIs the decoding accuracy during the ignored condition from that of the attended condition to develop a region-specific measure of attentional enhancement. We found that there was an enhancement in classification accuracy with attention across multiple cortical ROIs, particularly in the visual cortex, with some minor decrements in classification accuracy in frontal lobe ROIs (Fig 2c; difference maps showing statistical significance in Supplementary Figure 6a).

To quantify significant differences in classification performance across lobes and conditions, we computed mean classification performance across lobes for each condition (Fig 3a,c) and used a three-way repeated measures analysis of variance (rmANOVA), with factors for lobe (occipital/temporal/parietal/frontal) x hemisphere (left/right) x attention condition (attended/ignored) to test the significance of these results. The qualitative analysis of classification accuracy combined with the rmANOVA revealed three main findings: (i) category information varied by lobe (main effect of lobe: *F*(3,60)=49.76, *p*<.001, *η*_p_^2^=.713) with highest classification accuracy in the occipital lobe (Fig 3a,c blue bars, Tukey HSD: all *p*s <.001), (ii) category information varied across attended and ignored conditions (main effect of condition: *F*(1,20)=9.691, *p*=0.006, *η*_p_^2^=.326) with higher classification accuracy during the attended than ignored condition (Fig 3a,c dark vs. light bars, Tukey HSD: *p*<.001), and (iii) the effect of attention varied across lobes: significant interaction between lobe and condition (*F*(3,60)=29.080, *p*<.001, *η*_p_^2^=.593), whereby there was a larger difference between attended and ignored conditions in the occipital lobe and a smaller difference between conditions in the frontal lobe (Tukey HSD: *p*s <.001). There were no other significant effects (no main effect of hemisphere, interaction between condition and hemisphere, interaction between lobe and hemisphere, or three-way interaction between condition, hemisphere, and lobe; *p*s > .05). Results are similar when (i) examined separately for each category (faces, bodies, houses, and words; Supplementary Figure 7; Supplementary Table 3), suggesting that the results are not driven by a specific salient category, and (ii) when classification analyses of distributed responses were done for the raw signal amplitudes (Supplementary Figure 3).

**Figure 3.**
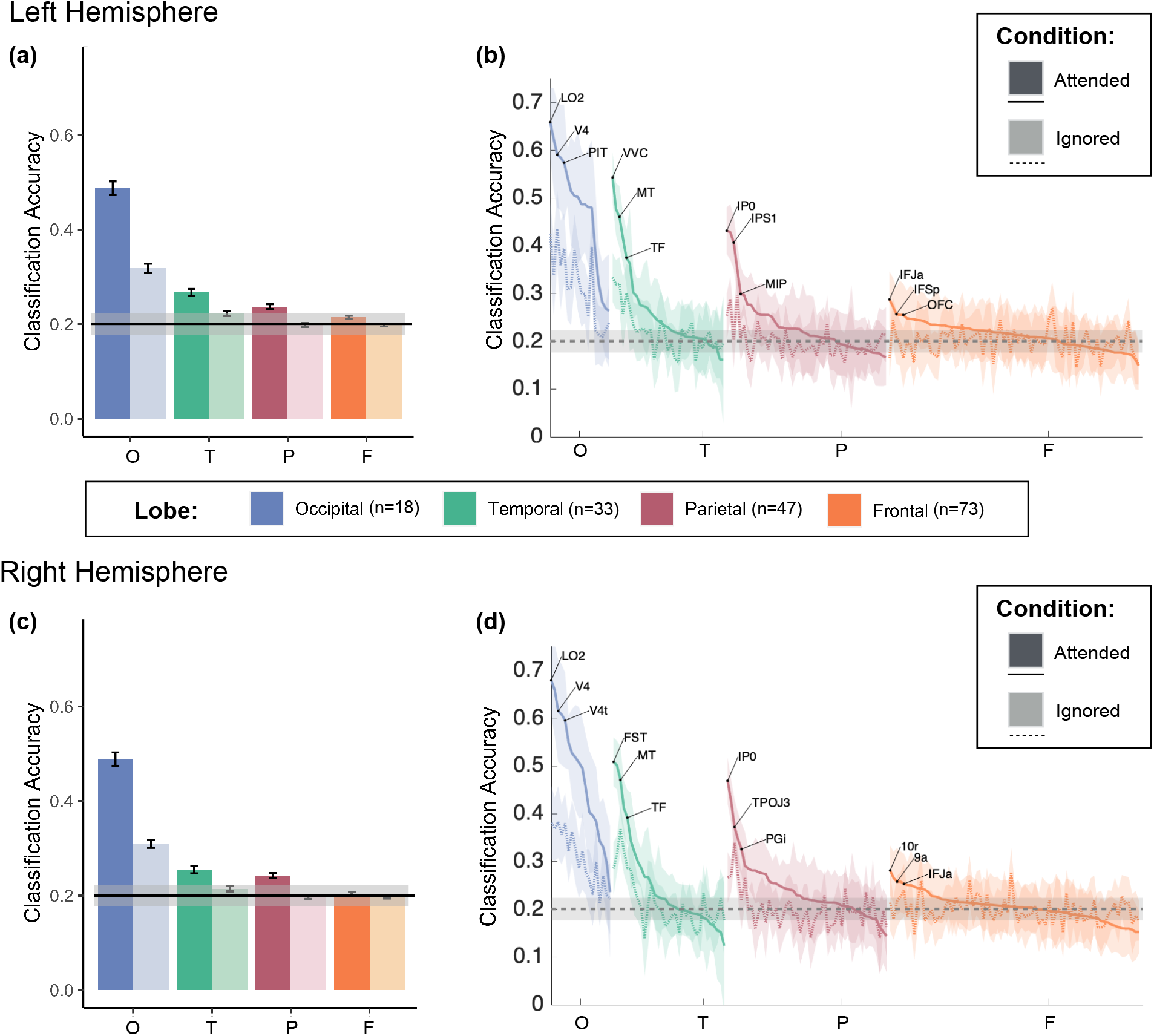
Classification accuracy by lobe, hemisphere, and condition. (a,c) Mean classification accuracy averaged across Glasser ROIs from each lobe: occipital (blue), temporal (green), parietal (red) and frontal (orange) lobes, in the left (a) hemisphere and right (c) hemisphere by condition: attended (dark colors) and ignored (light colors). *Error bars*: standard error of the mean across ROIs. (b,d) Same conventions as (a,c) but for each ROI separately. ROIs in the attended (solid line) and ignored (dashed line) conditions are ordered by the mean classification accuracy in the attended condition; *shaded area*: standard error of the mean across subjects. O: Occipital; T: Temporal; P: Parietal; F: Frontal. *Horizontal lines*: chance level; *shaded region*: the 95% confidence interval.

We further determined which Glasser ROIs received the greatest boost from attention by sorting the ROIs in each lobe by their classification performance in the attended condition and directly visualized classification accuracy across conditions for each ROI (Fig 3b,d). Results indicate that ROIs that have higher classification accuracy for the attended condition also tend to have higher classification accuracy for the ignored condition. Additionally, ROIs with higher classification accuracy during the ignored condition also have larger gains in classification performance during the attended condition than ROIs that have lower classification accuracy. To quantify this attentional enhancement, we ran a linear regression examining the relationship between mean decoding of category information for attended vs. ignored stimuli across ROIs, separately for each lobe and hemisphere. This approach allowed us to calculate a single scaling factor (the β value from the linear model) representing the attentional scaling factor for each lobe and hemisphere. Scaling significantly greater than 1 reflects attentional enhancement and scaling significantly less than 1 reflects attentional suppression. Results show that a linear model well captures the relationship between category information for attended vs ignored stimuli (all lobes and hemispheres *p*s < .05, Bonferroni corrected for multiple comparisons, except for the left frontal lobe *p*=.12, full stats in Table 1). The attentional scaling factor was 1.47 for the left and 2.02 for the right occipital lobe, 1.92 for the left and 1.60 for the right temporal lobe, 1.24 for the left and 1.14 for the right parietal lobe, indicating significant attentional enhancement of category representations bilaterally in these lobes. In the frontal lobe, the linear relationship was not significant after Bonferroni correction and showed a different trend in that the attentional scaling factor was less than 1 (0.17 in the left and 0.27 in the right frontal lobe). This indicates a trend in which category information in the frontal lobe was more decodable during the ignored condition than the attended condition.

### 3.2 What is the nature of residual correlations between different cortical regions and ventral temporal cortex category-selective ROIs?

Having determined that the decodability of category information varies with attention across cortex, we next investigated a potential correlate of this attentional enhancement: the strengthening of residual correlations between category-selective regions of VTC and other regions of the brain. These residual correlations, computed after regressing out stimulus-evoked BOLD responses, are thought to contain non-stimulus-driven or top-down attention signals that are not time-locked to stimulus events. One possibility is that residual correlations with a category-selective VTC fROI would be significant and positive only when that region’s preferred category is attended. This would suggest that regions that show positive correlations with category-selective fROIs are involved in directing attention to the relevant stimuli. Alternatively, finding significant residual correlations with the category-selective fROI when its preferred category is either attended or ignored would suggest that both attending and ignoring involve top-down control.

To distinguish between these hypotheses, we examined correlations in residual activity between each of the Glasser ROIs with four category-selective fROIs of VTC selective to different categories while attending or ignoring each region’s preferred category. To capture correlations in ongoing activity that were not time-locked to stimulus presentation, we subtracted the stimulus-evoked response in each voxel, and then measured the correlation between the mean residual activity of each region of the Glasser Atlas, and the mean residual activity in each of the category-selective fROIs of VTC (mFus-Faces, CoS-Houses, OTS-Bodies, OTS-Words) when subjects were either selectively attending or ignoring each category-selective region’s preferred category (Methods, section 2.7). We refer to these correlations between residual activities as ‘residual correlations’ and only consider Glasser ROIs that did not overlap with the VTC category-selective regions. We visualized for each Glasser ROI its mean residual correlation across all 4 VTC category-selective regions and participants (Fig 4 and Fig 5b,c). Additional visualizations and statistics by category are shown in Supplementary Figure 8 and Supplementary Table 4, respectively. We summarized data per lobe (Fig 5a,c), and noted that these correlation values are well within the ballpark of what would be expected based on prior work (Tompary et al., 2018). We then tested whether there are significant differences in residual correlations to VTC fROIs using a three-way repeated-measures ANOVA with factors of lobe (occipital/temporal/parietal/frontal) x hemisphere (left/right) x attention condition (attended/ignored). Results of these analyses reveal four main findings.

**Figure 4.**
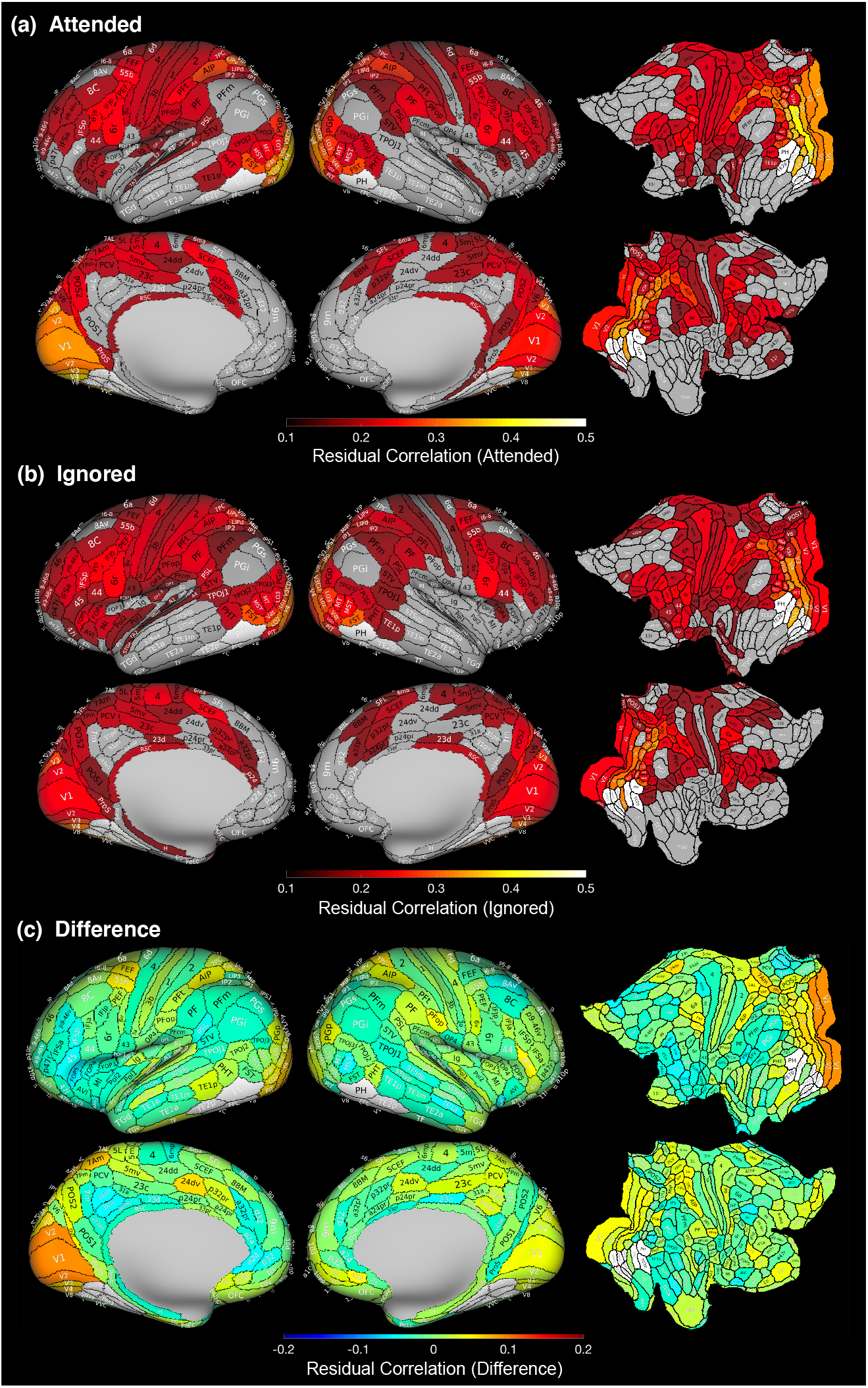
Comparisons of residual correlations with VTC category-selective regions during attended and ignored conditions. Mean correlations between residual activity in each Glasser ROI and residual activity in VTC category-selective regions, averaged across object categories and participants, in (a) the attended condition and (b) the ignored condition. Gray ROIs are those with residual correlations below 0.1. Colored ROIs are at least six standard deviations greater than chance level (0.01) for visualization purposes. (c) Mean difference between attended and ignored residual correlations. Visualization of these difference maps (c) depicting statistical significance for each ROI may be found in Supplementary Figure 6b.

**Figure 5.**
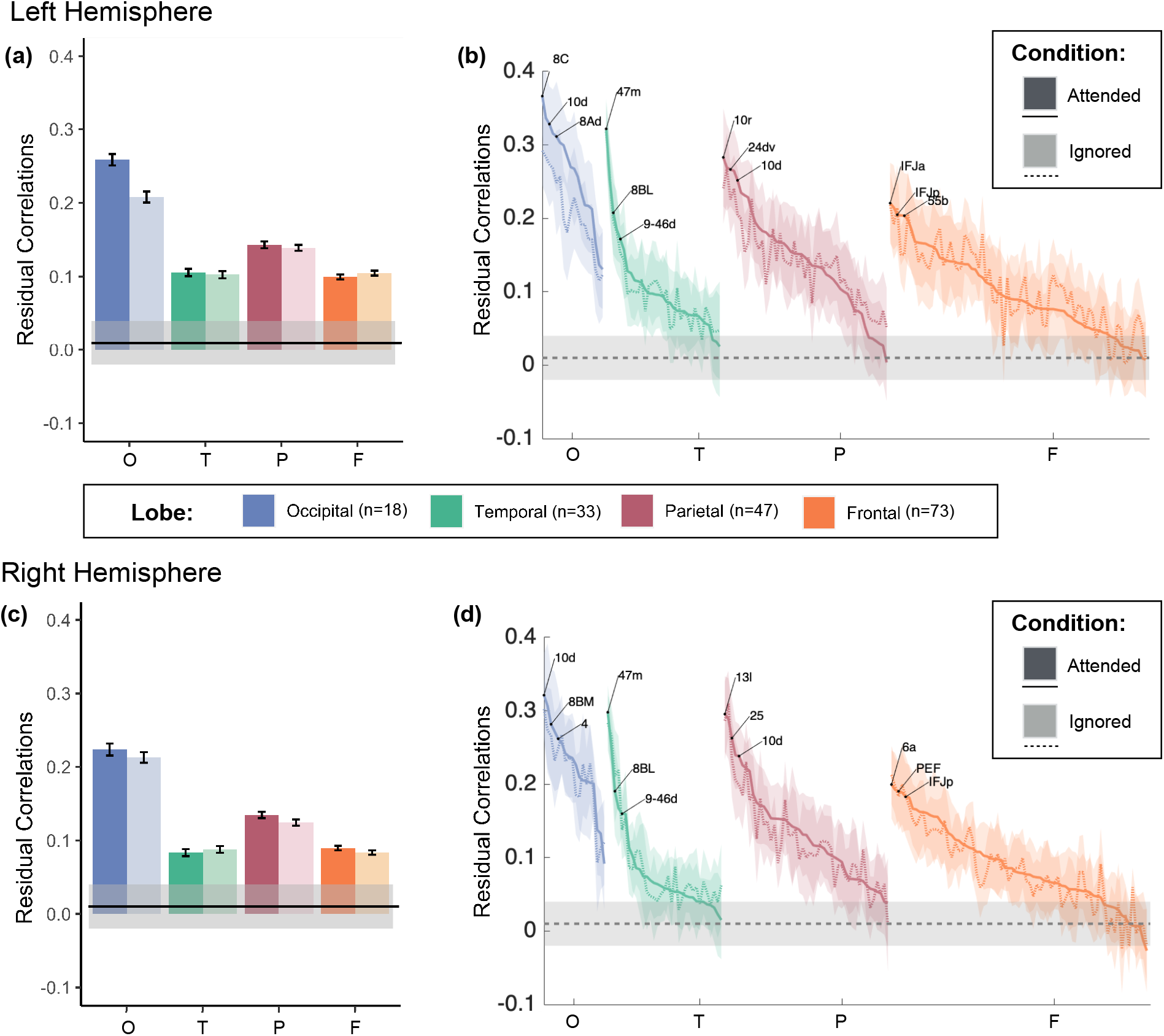
Residual correlations to VTC regions by lobe, hemisphere, and condition. (a,c) Mean residual correlations between VTC category-selective regions and each Glasser Atlas ROI from the occipital (blue), temporal (green), parietal (red) and frontal (orange) lobes, in the left hemisphere (a) and right hemisphere (c), respectively by condition: attended (dark colors) or ignored (light colors). *Error bars*: standard error of the mean across ROIs. (b,d) Same conventions as (a,c) but for each Glasser ROI separately. ROIs in the attended (solid line) and ignored (dashed line) conditions are ordered by the mean residual correlation in the attended condition; *shaded area*: standard error of the mean across subjects. O: Occipital; T: Temporal; P: Parietal; F: Frontal. *Horizontal lines*: chance level; *shaded region*: the 95% confidence interval.

First, as visible in Figs 4 and 5a,b, residual correlations to VTC fROIs are heterogeneous across brain lobes (main effect of lobe, *F*(3,60)=59.04, *p*<.001, *η*_p_^2^=.747). Surprisingly, we observed the highest residual correlations between ROIs in the occipital lobe (Fig 5a,c blue bars, Tukey HSD: *p*s <.001) despite having regressed out the stimulus-evoked hemodynamic response. This is also evident when examining individual Glasser ROIs, as residual correlations were above chance for all occipital ROIs (Fig 5b,d), but for only some of the ROIs in the other lobes. Second, we observed a significant interaction between lobe and attention condition (*F*(3,60)=4.96, *p*=0.004, *η*_p_^2^=.199), with the greatest difference between the attended and ignored conditions in the occipital lobe. The difference between the residual correlations per ROI for attended and ignored conditions is visualized on the Glasser Atlas (Fig 4c) again showing the largest enhancement in ROIs of the occipital lobe. Consistent with the lack of the main effect of attention condition, in many ROIs the differences in residual correlations are not significant at the ROI level (Supplementary Figure 6b). Third, we observed a significant three-way interaction among lobe, hemisphere, and condition (*F*(3,60)=3.12, *p*=0.033, *η*_p_^2^=.135). This three-way interaction appears to be driven primarily by the observation that in the occipital lobe, residual correlations in the attended condition were higher than in the ignored condition in the left hemisphere (Tukey HSD: *p*<.001) but not the right hemisphere (Tukey HSD: *p*>.05), whereas this pattern was not observed in any other lobe (Tukey HSD: *p*s>.05).

Ordering the ROIs by residual correlations in the attended condition in descending order reveals that ROIs with high residual correlations to VTC fROIs are evident across the occipital, parietal, and temporal lobes (Fig 5b,d). In the occipital lobe, the top regions were intermediate and high level visual areas: bilateral V3, V3CD, V4, V8 and LO1, left hemisphere V1, V2, V4t, LO2 and PIT; in the temporal lobe: bilateral VVC; and in the parietal lobe: bilateral IP0, IPS1, and MIP, and left hemisphere LIPv (Fig 5b,d). We also observed that several Glasser ROIs overlapping the fronto-parietal network (FPN): 55b, 7PC, 7AL, 7PL, AIP, FEF, IP0, LIPv, V7 and VIP were among the top 15% of ROIs with the highest enhancement of residual correlations with attention.

To test whether the effects of selective attention on residual correlations vary by the magnitude of these residual correlations, we ran a linear regression relating the mean residual correlations in the attended and ignored conditions across ROIs, separately for each lobe and hemisphere. In line with our hypothesis, regions with higher residual correlations overall also received the largest attentional modulation in all lobes and hemispheres (*p*s < .001, all surviving Bonferroni correction for multiple comparisons, full stats in Table 2). The attentional scaling factor was 1.18 for the left and 0.96 for the right occipital lobe, 0.98 for the left and 0.93 for the right temporal lobe, 1.25 for the left and 0.99 for the right parietal lobe, and 0.91 for the left and 0.91 for the right frontal lobe. The attentional scaling factors for the residual correlations are smaller in magnitude than those for category classification accuracy and reveal attentional enhancement (scaling larger than 1) only in the left occipital lobe and left parietal lobe, with the largest attentional suppression (scaling less than 1) in the bilateral frontal lobes.

### 3.3 Does the category decodability within a given region correlate with the strength of residual correlations between that region and VTC?

Our results thus far demonstrate that there is variability in the extent to which particular ROIs contain visual category information, as well as variability in the strength of their residual correlations with VTC category-selective fROIs. Thus, we asked: (1) is there a significant correlation between classification accuracy and the strength of residual correlations with VTC fROIs? (2) Does this relationship vary across attended and ignored conditions? We reasoned that finding a positive correlation between these metrics as well as a higher correlation among these metrics during attended than ignored conditions may suggest that attention plays an active role in enhancing task relevant category information by means of correlated activity among ROIs. To test this hypothesis, we measured the correlation between mean category classification accuracy and residual correlations with VTC fROIs. This analysis was done across the Glasser ROIs in each lobe (excluding the ROIs that overlap the VTC fROIs) and separately for the attended and ignored conditions.

Results shown in Fig 6 reveal three main findings. First, we found significant correlations between classification accuracy and residual correlation to category selective ROIs in the occipital, temporal, and parietal lobes (Fig 6; full statistics in Supplementary Table 5). That is, regions that show higher classification accuracy also have higher residual correlations with category-selective regions of VTC. Second, these correlations varied across lobes. In particular, these correlations were higher in the occipital and temporal lobes, and lower in the parietal and frontal lobe. Third, these associations varied by attention condition. For each hemisphere and attention condition, a step-wise linear regression model relating classification accuracy to residual correlations revealed significant improvement in model variance explained with the addition of attention condition as a factor (*p*s<.001) and with the addition of an interaction term between residual correlations and attention condition (*p*s<.001). Table 3 summarizes the results of these linear regressions including the interaction term by lobe. These interactions were significant, indicating a strengthening of the relationship between residual correlations and classification accuracy with attention, in the bilateral temporal lobes and the right parietal lobe, although the latter did not pass correction for multiple comparisons.

**Figure 6:**
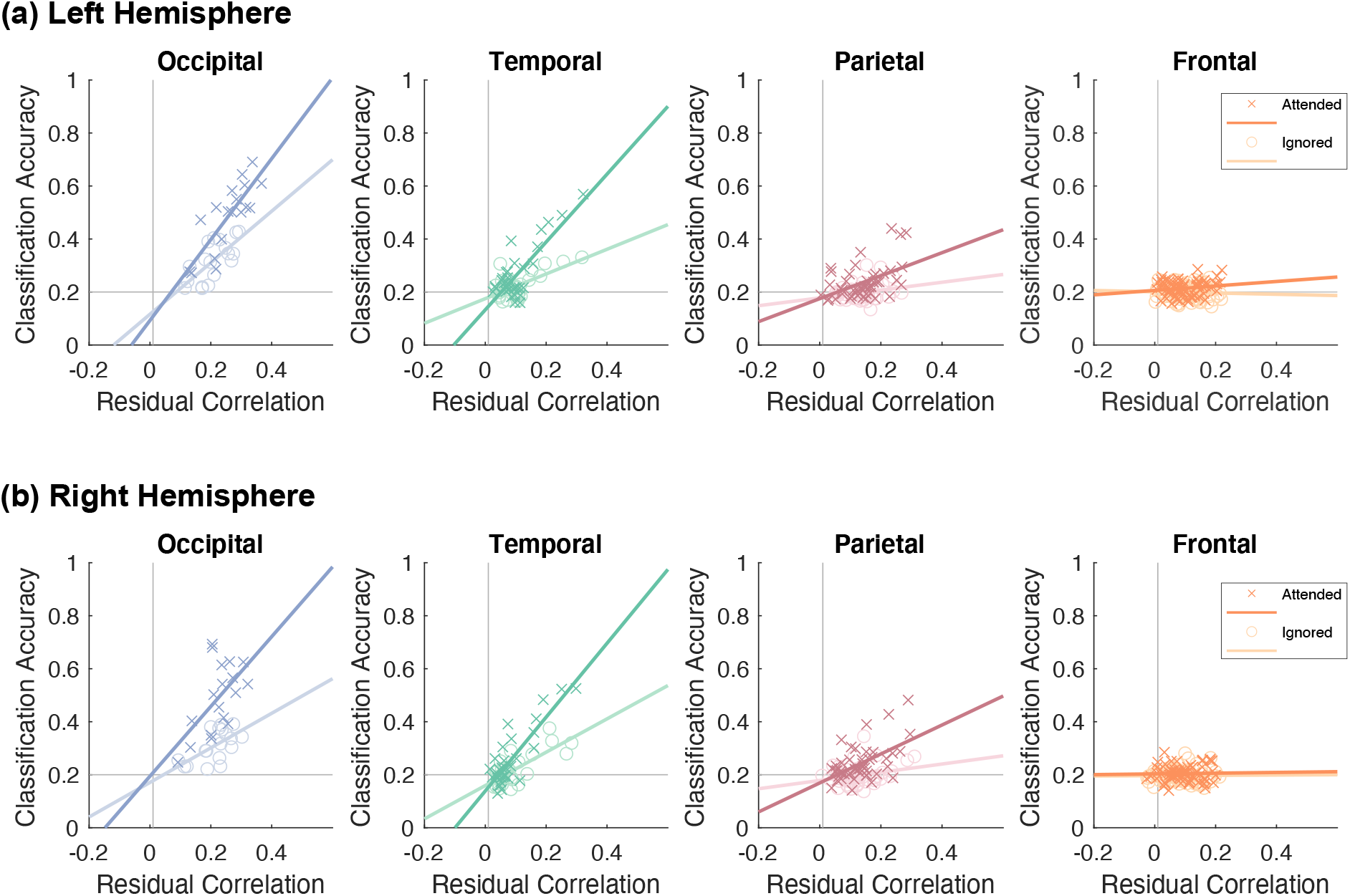
Attention enhances the correlation between classification accuracy and residual correlations. Each scatterplot depicts the relationship between mean classification accuracy (y-axis) and mean residual correlations (x-axis). Each point is a Glasser Atlas ROI marked by condition (dark-colored X’s: attended; light colored O’s: ignored). Data are shown separately for the (a) left hemisphere and (b) right hemisphere. Panels are arranged by lobe: *blue*: occipital; *green*: temporal; *red*: parietal; *orange*: frontal.

**Table 3:**
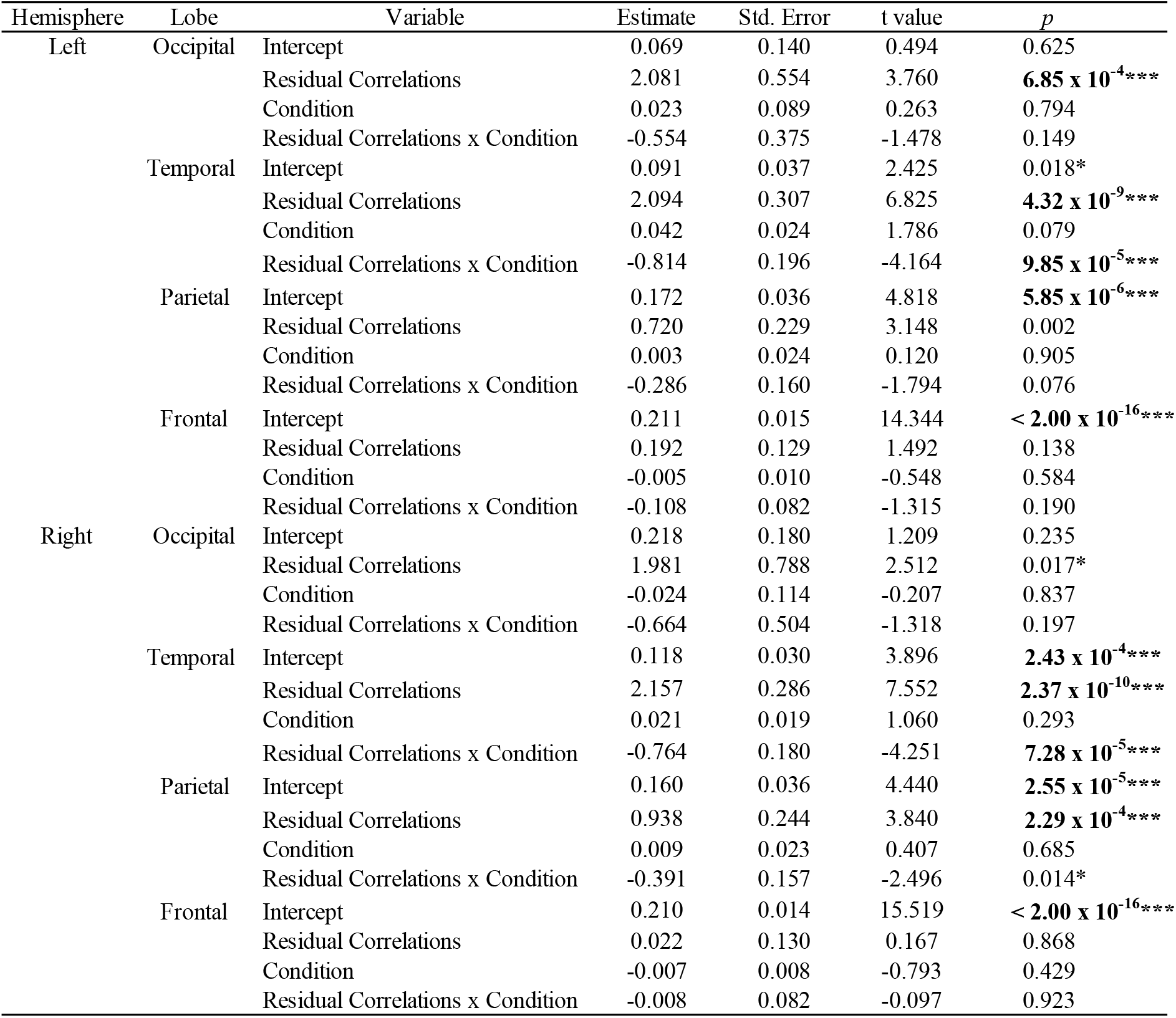
Linear models assessing the relationship between classification accuracy and residual correlations in attended and ignored conditions. Residual correlations in the attended and ignored conditions were used to predict classification accuracy values in each lobe and hemisphere. Bold values represent *p*-values surviving multiple comparisons correction (Bonferroni, 8 linear models). \**p*<.05, \*\**p*<.01, \*\*\**p*<.001.

Since the FPN has been hypothesized to be involved in top-down modulation, we also computed linear regressions between classification accuracy and residual correlations for the Glasser Atlas ROIs overlapping the FPN (Supplementary Figure 9), which are a subset of the frontal and parietal ROIs. This analysis revealed significant interactions between residual correlations and attention condition in both the left and right hemisphere fronto-parietal network (*p*s<.05, Bonferroni corrected; Supplementary Figure 9). As a comparison, we computed these relationships using the subset of frontal and parietal ROIs after excluding ROIs overlapping the FPN and found that these interactions were lower than those for FPN ROIs and failed to reach statistical significance after correction for multiple comparisons with the exception of the right frontal lobe (Supplementary Figure 10).

Together, these analyses show that there is a strong relationship between residual correlations and classification accuracy in the occipital, temporal, and parietal lobes, with significant attentional modulation in the bilateral temporal lobes. This suggests that regions spanning the occipital and parietal lobes show coupling between category information and residual correlations independent of attention condition while temporal lobes show enhanced coupling when stimuli are attended and category information is task-relevant. This effect appears to be primarily driven by increased category information when stimuli of these categories are attended.

## 4. Discussion

In this study, we investigated the ways that selective attention can influence visual category classification accuracy and the strength of residual correlations with VTC category-selective ROIs across the human cortex, and then assessed the relationship between these measurements. Our data reveal three main findings. First, we found that when two objects are simultaneously viewed, the category of the attended object can be decoded more readily from distributed responses in an ROI than the category of the ignored object. Second, we found that the strength of residual correlations with category-selective regions of VTC was higher when those regions’ preferred categories were attended compared to when they were ignored. Third, we found a positive correlation between classification accuracy and the strength of residual correlations to VTC category-selective ROIs, indicating that the stronger the residual correlations in a given region of cortex, the better we could decode category information from that region. Below we discuss the implications of these findings on elucidating the neural mechanisms of selective attention.

### Classification Accuracy of Attended and Ignored Object Category Representations

Our data contribute to the long-standing debate about whether attentional selection occurs “early” (Gandhi et al., 1999; Martínez et al., 1999; Somers et al., 1999) or “late” (Seidl et al., 2012; Shomstein et al., 2019; Wojciulik and Kanwisher, 1999) in the visual processing hierarchy (Yantis and Johnston, 1990). Prior evidence shows that attention impacts neural activity in visual cortex in many ways, including increasing neural firing rates (Motter, 1993) and tuning neural responses (Desimone and Duncan, 1995; Kastner et al., 1999, 1998; Reynolds and Heeger, 2009). Other evidence from stroke patients with damage to the right parietal cortex (a condition known as hemi-spatial neglect) reveal that information about visual objects that are outside the scope of attention still traverses quite far in the brain without being fully suppressed. For example, a hemi-neglect patient who was unable to attend to the left visual field was still capable of avoiding danger signals on the left side of a cartoon house (Marshall and Halligan, 1988). These data suggest that information about ignored visual objects may still be present in the brain even if this information is not accessible to conscious awareness. As such, it is noteworthy that we found greater enhancement of classification accuracy with attention not in the temporal lobe, but rather, in early visual and intermediate regions in the occipital lobe, which are thought to encode low-level and mid-level visual features. This finding is in line with predictions of the Reverse Hierarchy Theory (Hochstein and Ahissar, 2002) which suggests that focal attention to objects enhances low-level visual features relevant to the task of identifying objects rather than the category, or gist, which is encoded in high-level visual regions of VTC.

Our data are also consistent with the large body of research suggesting that attended stimuli can be decoded across many regions of the human brain (Kay et al., 2015; Bugatus et al., 2017; Çukur et al., 2013; Lee Masson et al., 2016; Peelen et al., 2009; Weiner and Grill-Spector, 2010; Klein et al., 2014). This includes studies using similar superimposed semi-transparent stimuli to demonstrate the effect of object-based attention in face- and house-selective regions of high-level visual cortex (O’Craven et al., 1999; Serences et al., 2004). It is possible that if we had used a different attention task, such as a spatial attention task, we may have found larger attentional effects in the parietal lobe, in line with previous spatial attention results (Peelen & Kastner, 2009; Sprague & Serences, 2013). These results also extend prior findings from our lab showing that both attended and unattended category information can be decoded from distributed responses across the entire lateral occipitotemporal complex (LOTC) as well as ventral temporal cortex (VTC), but that only attended information can be decoded from distributed responses across the entire ventrolateral prefrontal cortex (VLPFC) (Bugatus et al., 2017). Extending our prior results, we not only show the effects of selective attention to attended vs ignored stimuli relative to training with independent data using a different (oddball) task but also reveal the effect of attention at a finer spatial resolution across the entire brain. One potential mechanism that may underlie improved classification accuracy to attended vs ignored objects may be increased amplitude of BOLD activity and consequently higher signal-to-noise ratio (Buracas and Boynton, 2007; Gandhi et al., 1999; Martínez et al., 1999; Somers et al., 1999; Wojciulik and Kanwisher, 1999). Future studies would be needed to address whether this association is causal.

Our data also suggest that attention may function both by enhancing sensory representations in visual cortex (Baldauf and Desimone, 2014; Cohen and Tong, 2015; Zhou et al., 2015), and by flexibly altering the readout of those sensory representations in higher order cortical areas (Birman and Gardner, 2019; Bugatus et al., 2017; Peelen et al., 2009). Although we observed significant enhancement in classification accuracy with attention, we found that both ignored and attended categories could be classified significantly above chance in visual cortex. This implies that if the readout from the visual cortex to the frontal cortex were fixed, then category information would be decodable in the frontal lobe in both attended and ignored conditions, as in visual cortex. However, contrary to this prediction, we found that category information in the frontal lobe is only decodable for attended stimuli. This suggests a flexible readout of task-relevant information in the frontal lobe and that attention may enable the transfer of information from visual cortex to the frontal lobe.

### Residual Correlations May Reflect Sharing of Attended Information Between Cortical Regions

Two aspects of our data support the hypothesis that residual correlations may reflect the sharing of information about attended visual object categories between sensory regions and higher-order cortices. First, we found that occipital and temporal lobes had both the strongest category information and the strongest coupling between classification accuracy and residual correlations. Second, we found that the bilateral temporal lobes showed the greatest enhancement of coupling between classification accuracy and residual correlations with attention. Future studies using causal manipulations of information transfer in the brain may further illuminate whether these residual correlations play an active role as well as have behavioral consequences.

Another interesting observation is that there were still substantial residual correlations between brain regions and category-selective regions in VTC when those regions’ preferred categories were ignored. This suggests that top-down control may be necessary not only when attending stimuli but also when ignoring stimuli, in line with prior work emphasizing the importance of top-down control for both enhancement and suppression of sensory information (Martinez-Trujillo and Treue, 2004; Scolari et al., 2012). While we acknowledge that our residual correlations do not inform about the directionality of information flow, our results set the stage for future studies to assess the causal relationships between category information and residual correlations. For example, future studies could test whether manipulation of residual correlations between sensory and higher-order cortices directly affects the availability of category information in either of these regions.

### Implications for Clinical Research

Attention difficulties are among the most debilitating symptoms of mental disorders (Cotrena et al., 2016; Fehnel et al., 2013) and are associated with poorer prognoses (Majer et al., 2004), yet they are often overlooked. In fact, they remain among the least well-understood neurobiologically (Keller et al., 2019b) outside of well-documented attentional biases toward negative information (Joorman and Gotlib, 2010). Recent work has shown that visual selective attention in particular is severely impaired in a subset of individuals with Major Depressive Disorder (Keller et al., 2019a) and individuals with symptoms of generalized/physiological anxiety (Keller et al., 2021). In line with burgeoning efforts in psychiatric research to understand transdiagnostic dimensions of psychopathology across units of analysis, known as the “RDoC” initiative (Insel et al., 2010), our work advances our understanding of the widespread cortical regions involved in selective attention, providing a roadmap for future studies to probe causal mechanisms in psychiatric populations.

Two key challenges in addressing selective attention impairments in depression and other mental illnesses are: (1) the observation that selective attention impairments in depression are often not alleviated with current first-line antidepressant pharmacotherapy (Keller et al., 2019c), and (2) the lack of precise neural targets for novel treatment development targeting specific symptom dimensions (Williams, 2016). First, to reduce the burden on patients to undergo multiple rounds of treatment attempts (often with debilitating side effects) before finding an effective treatment, future studies may utilize our behavioral paradigm to develop a clinic-ready measure of attention impairment for guiding more personalized treatment selection among currently-available options. Second, our analysis of visual selective attention utilized a data-driven whole-brain approach to look beyond sensory cortices, opening the door for the potential development of stimulation therapies targeting a wider range of accessible brain areas. Thus, our study of attention using a low-cost behavioral paradigm lends itself to translational efforts for mapping attentional difficulties in various psychiatric populations.

## Conclusions

Our study demonstrates that correlations in residual activity between higher-order brain areas and the ventral temporal cortex is related to the sharing of task-relevant object category information across cortical regions. Both decodability of object categories and residual correlations with regions preferentially processing these object categories are enhanced when said categories are attended compared to when they are ignored. Importantly, these findings inform our understanding of how selective attention influences the representation of information across the brain by revealing residual correlations between regions that may reflect the preferential sharing of attended information. Future studies may probe the directionality of this information flow using causal manipulations, which may have important implications for clinical research on selective attention impairments in psychiatric illness.

## 5. Acknowledgments

We thank John Cocjin for fruitful discussions and for contributions to an earlier version of this study.

## 6. Funding Sources

This work was supported by the National Eye institute (NEI) grant R01EY023915 to KGS. ASK. is supported by the National Defense Science and Engineering Graduate Fellowship (NDSEG) and both ASK, AJ and LB received support from the Stanford Center for Mind, Brain, Computation and Technology (MBCT) Traineeship. This work was also supported by the National Institutes of Health [grant number U01MH109985 under PAR-14-281].

## 7. Declarations of Interest

LMW has received advisory board fees from Laureatte Institute of Brain Research and Psyberguide of the One Mind Institute, for work unrelated to this manuscript. The other authors declare no conflicts of interest.

## 8. Author Contributions

**Arielle S. Keller:** Conceptualization, Formal analysis, Methodology, Visualization, Software, Writing – original draft preparation, Writing – reviewing & editing.

**Akshay V. Jagadeesh:** Conceptualization, Formal Analysis, Methodology, Visualization, Software, Writing – reviewing and editing.

**Lior Bugatus:** Conceptualization, Data curation, Investigation, Writing – reviewing and editing.

**Leanne M. Williams:** Resources, Writing – reviewing and editing, Supervision.

**Kalanit Grill-Spector:** Conceptualization, Resources, Writing – reviewing and editing, Supervision.

