## Supplementary Material for "Attention enhances category representations across the brain with strengthened residual correlations to ventral temporal cortex"

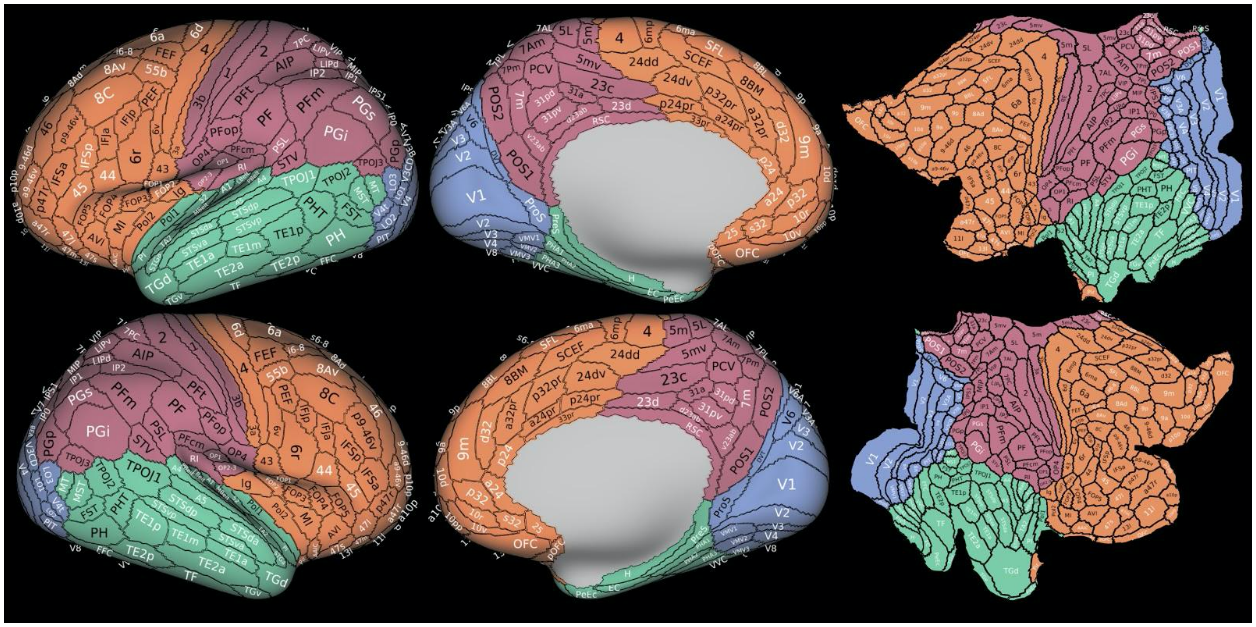


**Supplementary Figure 1. Glasser Atlas lobe designations**. *Orange:* frontal lobe; *red:* parietal lobe; *green:* temporal lobe; *blue:* occipital lobe.

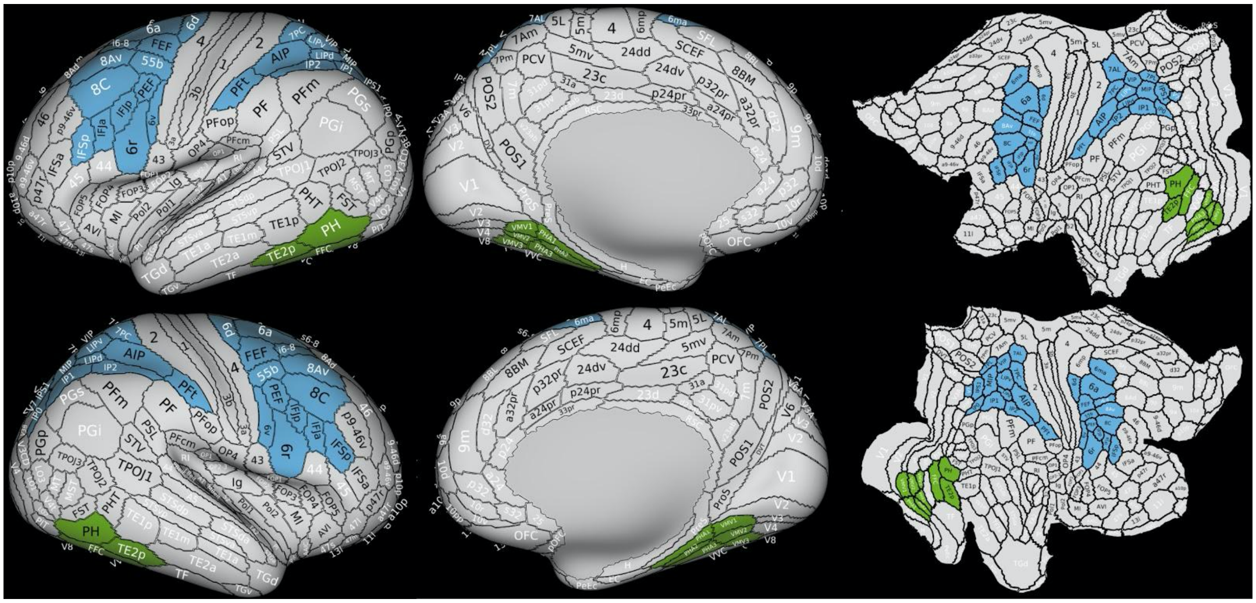


**Supplementary Figure 2. Glasser Atlas fronto-parietal network (FPN) and category-selective region designations.** *Blue:* regions overlapping the FPN (Osher et al., 2019); *green:* regions overlapping category-selective functional ROIs in the ventral temporal cortex.

| Subject | Left Hemisphere | | | | Right Hemisphere | | | |
|---|---|---|---|---|---|---|---|---|
| | mFus-Faces | CoS-Houses | OTS-Bodies | pOTS-Chars | mFus-Faces | CoS-Houses | OTS-Bodies | pOTS-Chars |
| 1 | ✓ | ✓ | ✓ | ✓ | ✓ | ✓ | ✓ | ✓ |
| 2 | ✓ | ✓ | ✓ | | ✓ | ✓ | ✓ | ✓ |
| 3 | ✓ | ✓ | | ✓ | ✓ | ✓ | ✓ | ✓ |
| 4 | ✓ | ✓ | | ✓ | ✓ | ✓ | | ✓ |
| 5 | ✓ | ✓ | ✓ | ✓ | ✓ | ✓ | ✓ | ✓ |
| 6 | ✓ | ✓ | ✓ | ✓ | ✓ | ✓ | ✓ | ✓ |
| 7 | ✓ | ✓ | | ✓ | ✓ | ✓ | ✓ | ✓ |
| 8 | ✓ | ✓ | ✓ | ✓ | ✓ | ✓ | ✓ | ✓ |
| 9 | ✓ | ✓ | | ✓ | ✓ | ✓ | ✓ | |
| 10 | ✓ | ✓ | | ✓ | ✓ | ✓ | ✓ | ✓ |
| 11 | ✓ | ✓ | ✓ | ✓ | ✓ | ✓ | ✓ | |
| 12 | ✓ | ✓ | ✓ | ✓ | ✓ | ✓ | ✓ | ✓ |
| 13 | | ✓ | | ✓ | ✓ | ✓ | ✓ | ✓ |
| 14 | | ✓ | | ✓ | ✓ | ✓ | | ✓ |
| 15 | ✓ | ✓ | ✓ | | ✓ | ✓ | ✓ | |
| 16 | | ✓ | | ✓ | ✓ | ✓ | ✓ | ✓ |
| 17 | | ✓ | | ✓ | | ✓ | ✓ | |
| 18 | | ✓ | ✓ | | ✓ | ✓ | ✓ | |
| 19 | ✓ | ✓ | ✓ | | ✓ | ✓ | ✓ | |
| 20 | | ✓ | | ✓ | ✓ | ✓ | ✓ | |
| 21 | | ✓ | | ✓ | | ✓ | ✓ | |
| Total: | 14 | 21 | 10 | 17 | 19 | 21 | 19 | 13 |

**Supplementary Table 1.** Presence or absence of each functional region of interest (fROI) in ventral temporal cortex (VTC) by subject.

| Measurement | Hemisphere | Condition | r | *p* |
|---|---|---|---|---|
| Classification Accuracy | Left | Attended | 0.032 | 0.667 |
| | | Ignored | -0.070 | 0.351 |
| | Right | Attended | -0.022 | 0.768 |
| | | Ignored | -0.081 | 0.280 |
| Residual Correlations | Left | Attended | 0.117 | 0.119 |
| | | Ignored | 0.070 | 0.349 |
| | Right | Attended | 0.061 | 0.420 |
| | | Ignored | 0.030 | 0.690 |

**Supplementary Table 2.** Correlation between each of our key measurements (Classification Accuracy and Residual Correlations) with ROI size across all ROIs of the Glasser atlas. None of these correlations are significant.

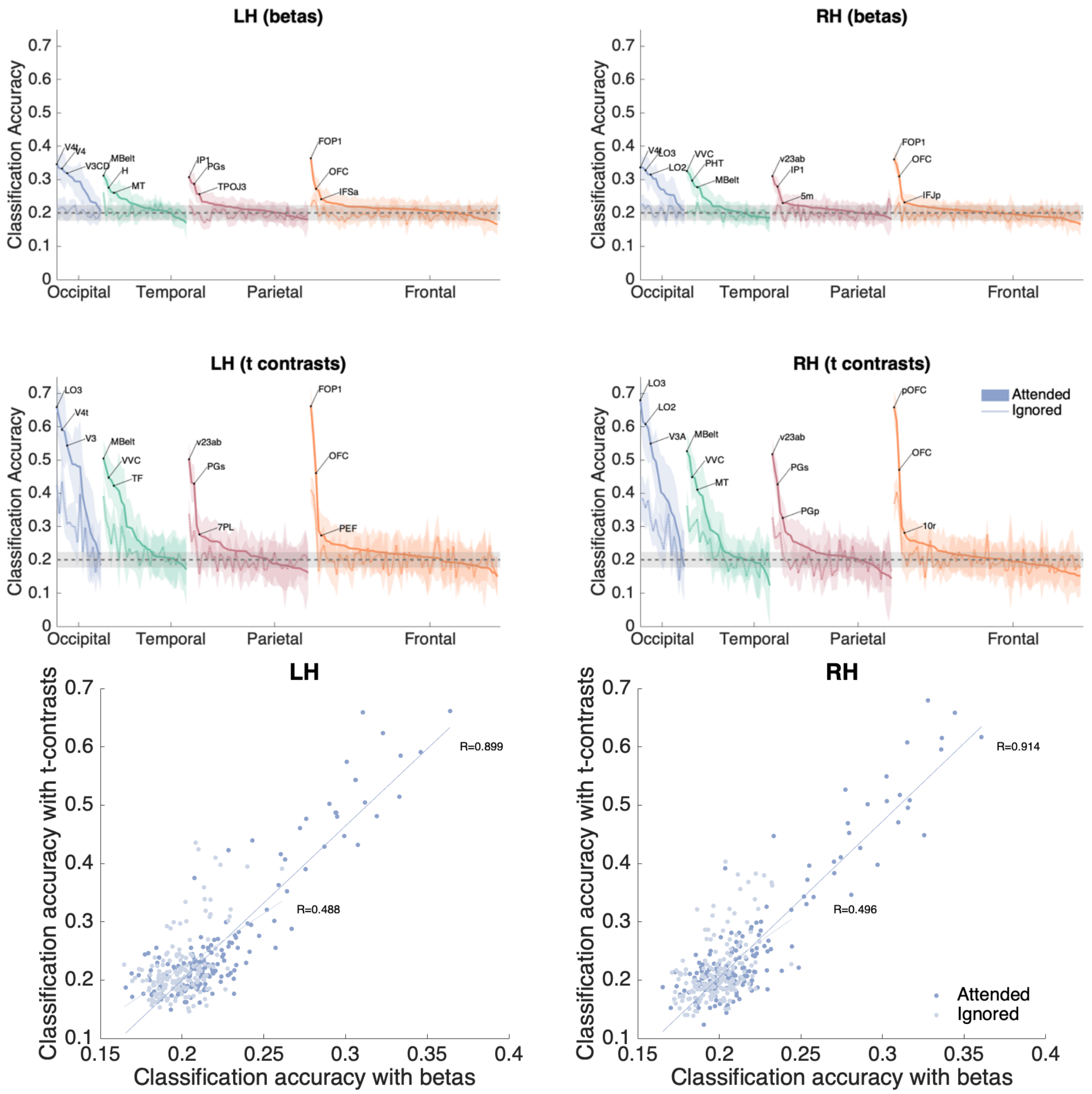


**Supplementary Figure 3. Comparison of classification results based on raw responses (beta values) vs. contrasts (t-values).** *Top row:* classification accuracy using beta values of each ROI, grouped by lobe, for left hemisphere (left) and right hemisphere (right). *Middle row:* classification accuracy using t-contrasts of each ROI, grouped by lobe, for left hemisphere (left) and right hemisphere (right). *Bottom row:* correlation between classification results obtained using t-contrasts and beta values for attended (dark blue) and ignored (light blue). Although it is the case that the classification accuracies are generally lower when using betas instead of t-contrasts, we replicate the main findings that classification accuracy varies by lobe ($F(3,60)=36.79$, $p<.001$), attentional condition ($F(1,20)=7.108$, $p=0.015$) and hemisphere ($F(1,20)=4.957$, $p=0.038$), and the significant interaction of lobe and attentional condition ($F(3,60)=26.1$, $p<.001$). *LH:* Left Hemisphere; *RH:* Right Hemisphere.

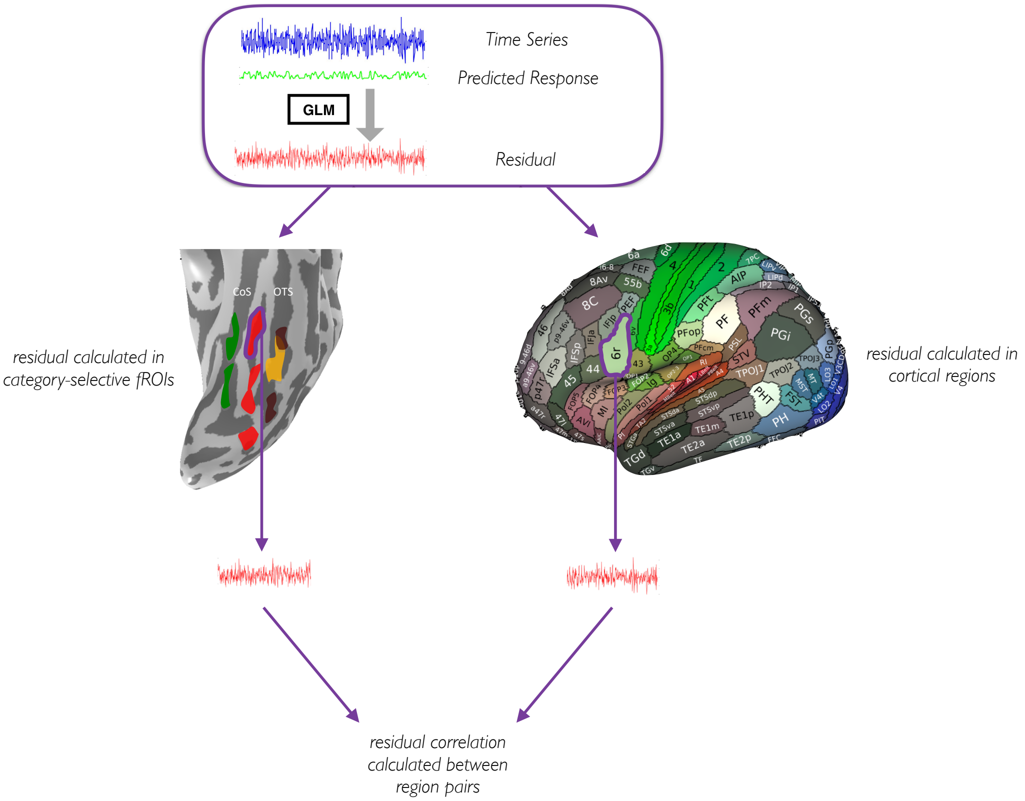


**Supplementary Figure 4. Residual correlations analysis pipeline (Methods Section 2.7).** We start with the time series of the BOLD response recorded from the fMRI scanner, then model the predicted hemodynamic response with a GLM, and extract the residual activity (unexplained variance) from this model. We compute residuals in four category-selective fROIs in VTC defined subject-by-subject with a functional localizer (left) as well as in all regions of the Glasser Atlas (Glasser et al., 2016). We then compute correlations between the time course of residual activity in each pair of regions (one cortical region with one category-selective fROI at a time), under two conditions: one in which that fROI's preferred category is attended and one in which the fROI's preferred category is ignored.

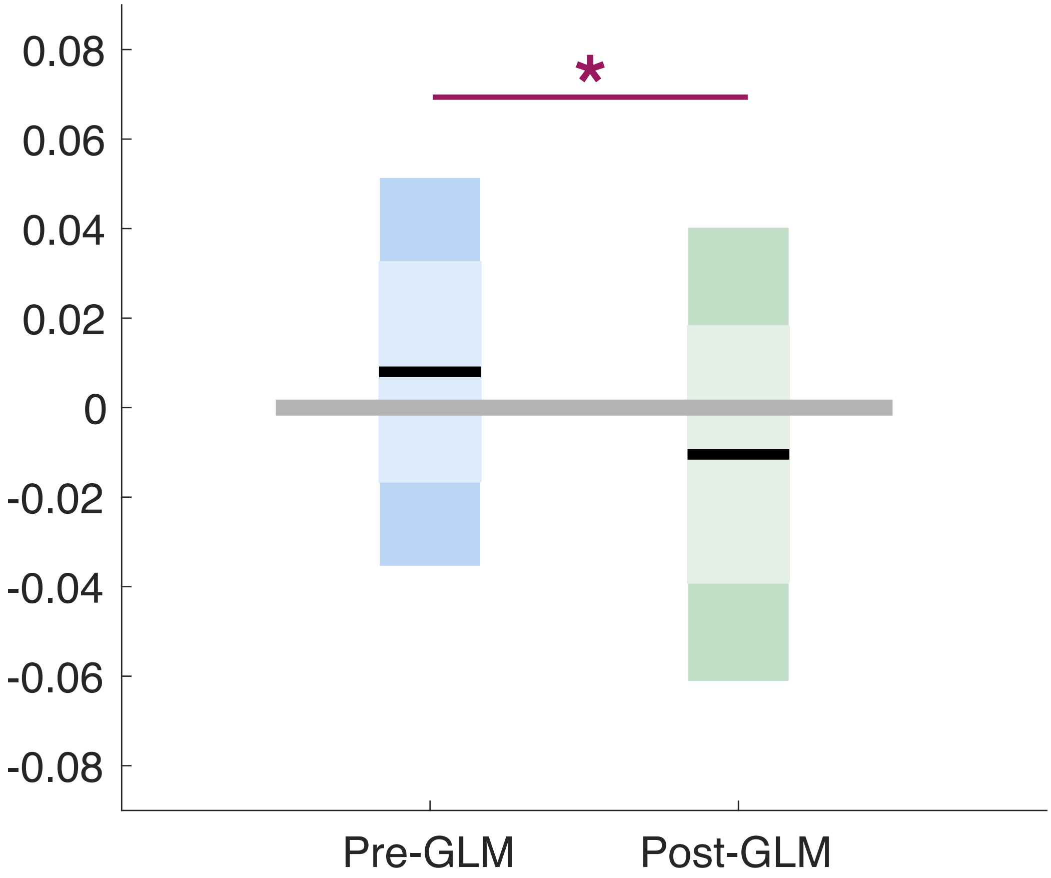


**Supplementary Figure 5. Control analysis validating the procedure for deriving the residual correlations.** To validate that the GLM captured the task-evoked activity, we computed correlations in the time series ("Pre-GLM") and residual activity ("Post-GLM") between two trials of the same condition averaged over each condition and each region of the Glasser atlas for each subject. Results show that the "Pre-GLM" distribution of correlations is significantly greater than the "Post-GLM" distribution, and the "Post-GLM" distribution was not significantly different from zero, validating our use of the GLM to remove consistent task-evoked activity across trials. $^{*}p<.05$.

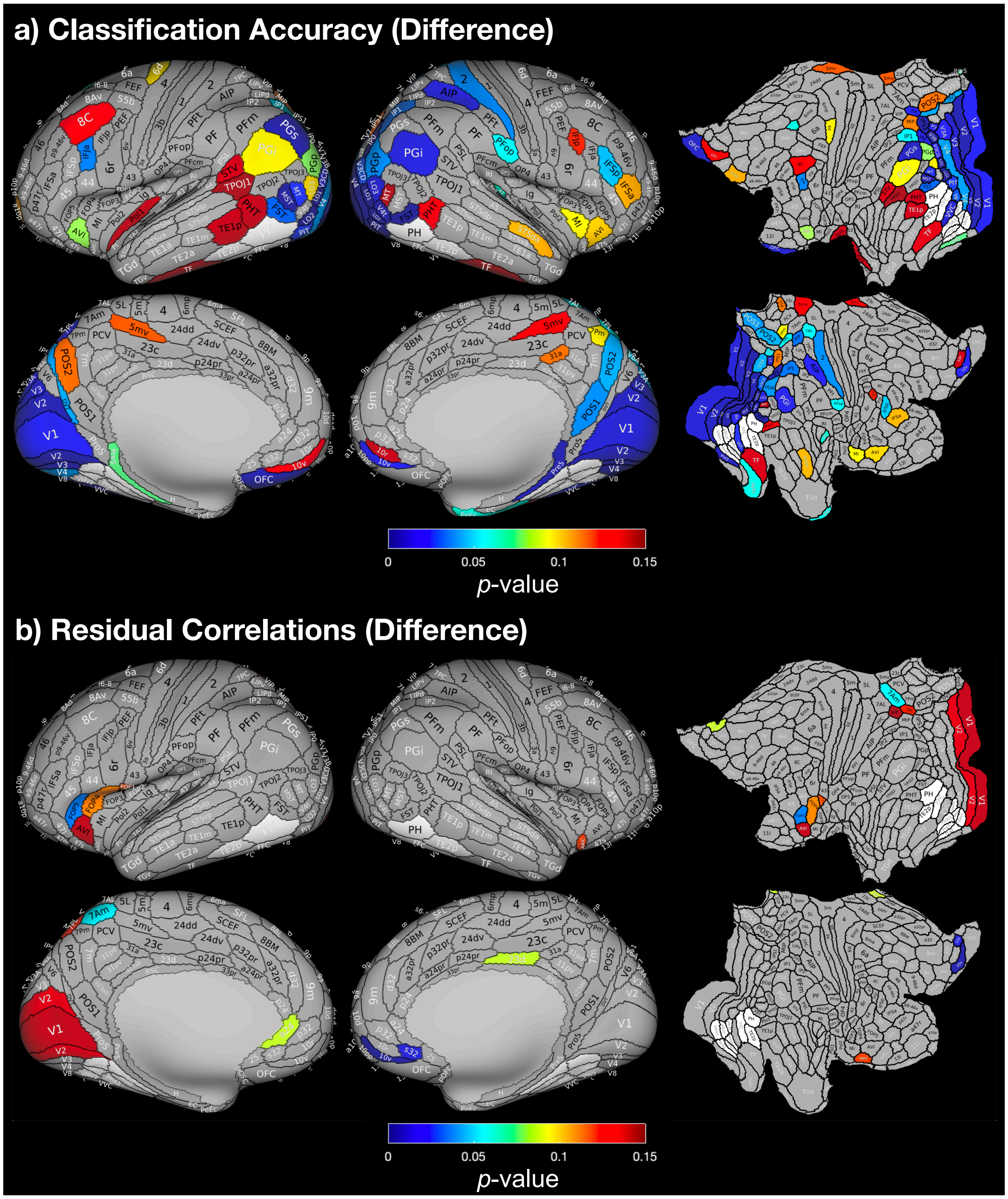


**Supplementary Figure 6. Brain maps depicting the significance (*p*-value) of the difference between attended and ignored conditions for the classification accuracy (a) and residual correlations (b).** To determine the statistical significance of the difference, we performed *t*-tests comparing the Attended to Ignored condition for each ROI in each hemisphere. Gray regions are those that did not pass a lenient *p*-value threshold of 0.15 (uncorrected) (lenient threshold used for visualization purposes only). Only three of the regions above (LO1, LO2 and PIT in the right hemisphere for the classification accuracy) remain significant after correction for multiple comparisons. Accordingly, all analyses focused on comparisons of lobe, hemisphere, and condition rather than differences between individual ROIs.

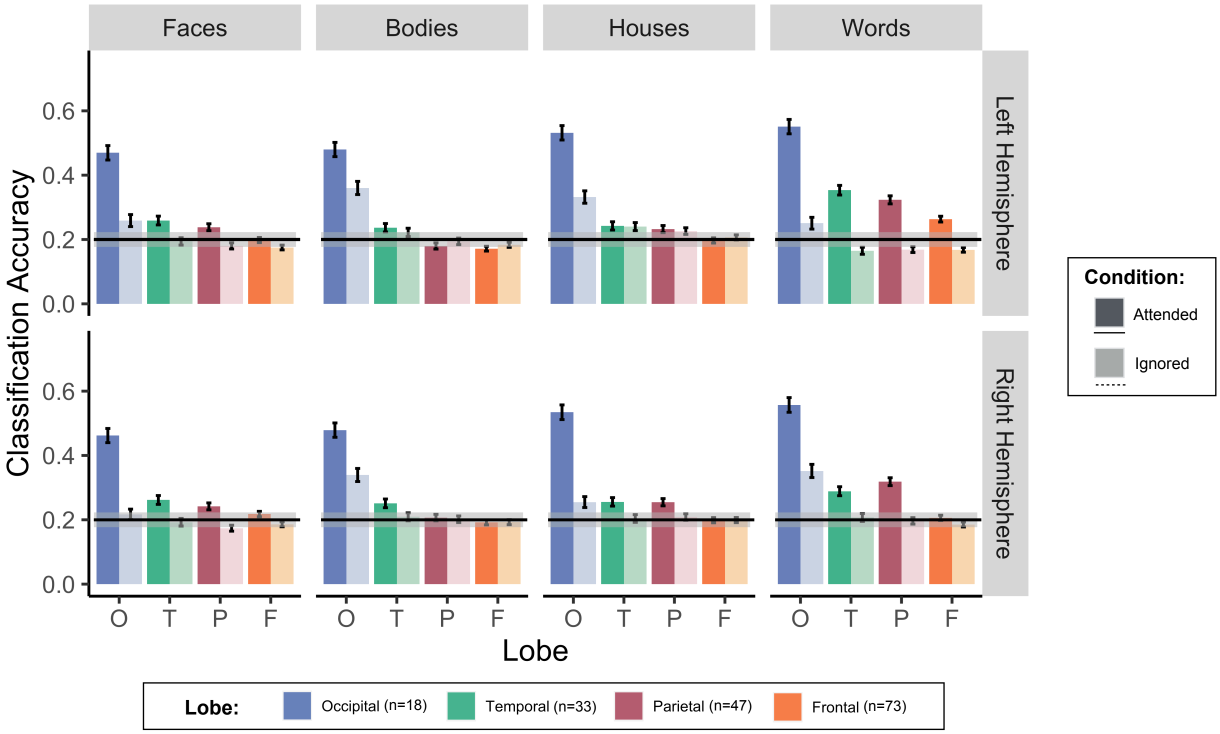


**Supplementary Figure 7. Classification accuracy by stimulus condition (faces, bodies, houses, words), hemisphere (left, right), lobe (occipital, temporal, parietal, frontal), and condition (attended, ignored).** *Error bars*: the standard error of the mean across ROIs in each lobe. *Horizontal lines:* chance level with an error cloud representing the 95% confidence interval. A three-way rmANOVA (Lobe x Condition x Stimulus Condition) revealed a non-significant main effect of stimulus condition ($F(3,60)=0.399$, $p=0.754$) but a significant interaction between stimulus condition and attended/ignored condition ($F(3,60)=5.529$, $p=0.002$). Note that the Cars condition was excluded because residual correlations are only calculated for stimulus conditions for which there is a corresponding category-selective fROI in VTC. Separate rmANOVAs by stimulus condition may be found in **Supplementary Table 3.**

| Stimulus Condition | ANOVA | Sum Sq | F value | *p* |
|---|---|---|---|---|
| Faces | Lobe | 15.47 | 16.21 | **$7.89 \times 10^{-8}$** |
| | Condition | 7.82 | 22.48 | **$1.25 \times 10^{-4}$** |
| | Lobe x Condition | 6.09 | 9.58 | **$2.92 \times 10^{-5}$** |
| Houses | Lobe | 33.66 | 24.07 | **$2.39 \times 10^{-10}$** |
| | Condition | 0.41 | 0.58 | 0.456 |
| | Lobe x Condition | 3.08 | 7.58 | **$2.22 \times 10^{-4}$** |
| Bodies | Lobe | 27.70 | 26.79 | **$4.04 \times 10^{-11}$** |
| | Condition | 2.26 | 2.56 | 0.125 |
| | Lobe x Condition | 9.19 | 19.23 | **$7.45 \times 10^{-9}$** |
| Words | Lobe | 29.91 | 21.57 | **$1.34 \times 10^{-9}$** |
| | Condition | 23.95 | 18.62 | **$3.36 \times 10^{-4}$** |
| | Lobe x Condition | 6.47 | 17.93 | **$2.00 \times 10^{-8}$** |

**Supplementary Table 3.** Results of the ANOVA on classification accuracy comparing lobe (occipital, temporal, parietal, frontal), and condition (attended, ignored) separately for each stimulus condition (faces, bodies, houses, words). Data were averaged over hemisphere to maximize the number of subjects included in each ANOVA. Bold values are those that pass Bonferroni correction for four comparisons (one ANOVA for each stimulus condition).

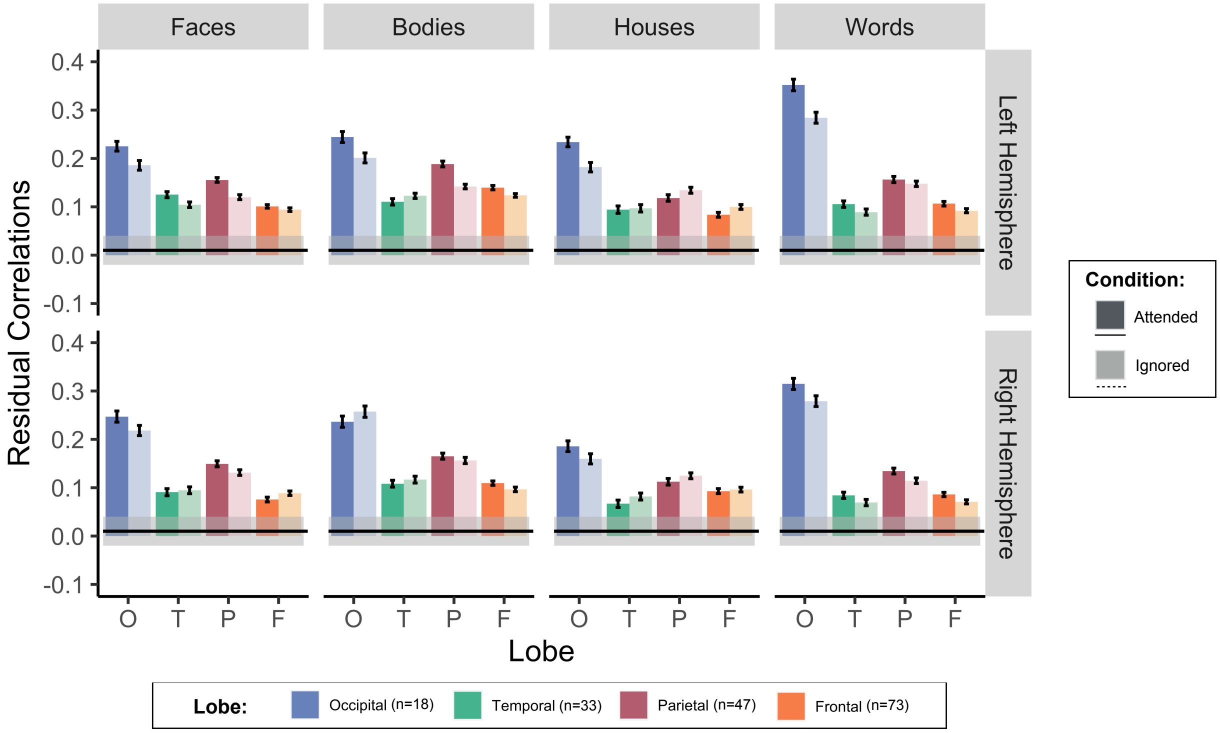


**Supplementary Figure 8. Residual correlations by stimulus condition (faces, bodies, houses, words), hemisphere (left, right), lobe (occipital, temporal, parietal, frontal), and condition (attended, ignored).** *Error bars:* the standard error of the mean across Glasser ROIs in each lobe. *Horizontal lines*: chance level with an error cloud representing the 95% confidence interval. The Cars condition was excluded because residual correlations were only calculated for stimulus conditions for which there is a corresponding category-selective fROI in VTC. Note also that we did not perform a three-way rmANOVA (Lobe x Condition x Stimulus Condition) on the residual correlations as we did for the classification accuracy given that some participants were missing individual VTC fROIs (separate rmANOVAs by stimulus condition may be found in Supplementary Table 4).

| Stimulus Condition | ANOVA | Sum Sq | F value | *p* |
|---|---|---|---|---|
| Faces | Lobe | 10.93 | 27.00 | **$8.37 \times 10^{-11}$** |
| | Condition | 0.07 | 0.26 | 0.614 |
| | Lobe x Condition | 0.30 | 3.18 | 0.031 |
| Houses | Lobe | 11.79 | 22.04 | **$1.33 \times 10^{-9}$** |
| | Condition | 0.09 | 0.09 | 0.772 |
| | Lobe x Condition | 0.21 | 1.56 | 0.210 |
| Bodies | Lobe | 6.94 | 19.04 | **$8.55 \times 10^{-9}$** |
| | Condition | 0.06 | 0.06 | 0.820 |
| | Lobe x Condition | 0.43 | 2.94 | 0.040 |
| Words | Lobe | 22.99 | 48.52 | **$5.74 \times 10^{-15}$** |
| | Condition | 0.45 | 1.25 | 0.279 |
| | Lobe x Condition | 0.24 | 1.56 | 0.211 |

**Supplementary Table 4.** Results of the ANOVA on residual correlations comparing lobe (occipital, temporal, parietal, frontal), and condition (attended, ignored) separately for each stimulus condition (faces, bodies, houses, words). Data were averaged over hemisphere to maximize the number of subjects included in each ANOVA. Bold values are those that pass Bonferroni correction for four comparisons (one ANOVA for each stimulus condition).

| Lobe | Hemisphere | N | Condition | R | *p* |
|---|---|---|---|---|---|
| Occipital | Left | 18 | Attended | 0.817 | **3.550 x $10^{-5}$*** ** |
| | | | Ignored | 0.709 | **0.001** ** |
| | Right | 18 | Attended | 0.587 | 0.010* |
| | | | Ignored | 0.622 | 0.006** |
| Temporal | Left | 33 | Attended | 0.809 | **1.180 x $10^{-8}$*** ** |
| | | | Ignored | 0.643 | **0.0001*** ** |
| | Right | 33 | Attended | 0.856 | **2.271 x $10^{-10}$*** ** |
| | | | Ignored | 0.756 | **3.708 x $10^{-7}$*** ** |
| Parietal | Left | 47 | Attended | 0.480 | **0.0006*** ** |
| | | | Ignored | 0.249 | 0.092 |
| | Right | 47 | Attended | 0.536 | **0.0001*** ** |
| | | | Ignored | 0.257 | 0.082 |
| Frontal | Left | 73 | Attended | 0.166 | 0.160 |
| | | | Ignored | -0.050 | 0.676 |
| | Right | 73 | Attended | -0.027 | 0.822 |
| | | | Ignored | 0.013 | 0.914 |

**Supplementary Table 5. Correlations between classification accuracy and residual correlations in attended and ignored conditions.** Correlations in the attended and ignored conditions were evaluated in each lobe and hemisphere. Bold values represent correlations with *p*-values surviving multiple comparisons correction (Bonferroni, 16 comparisons). *$p<.05$, **$p<.01$, ***$p<.001$.

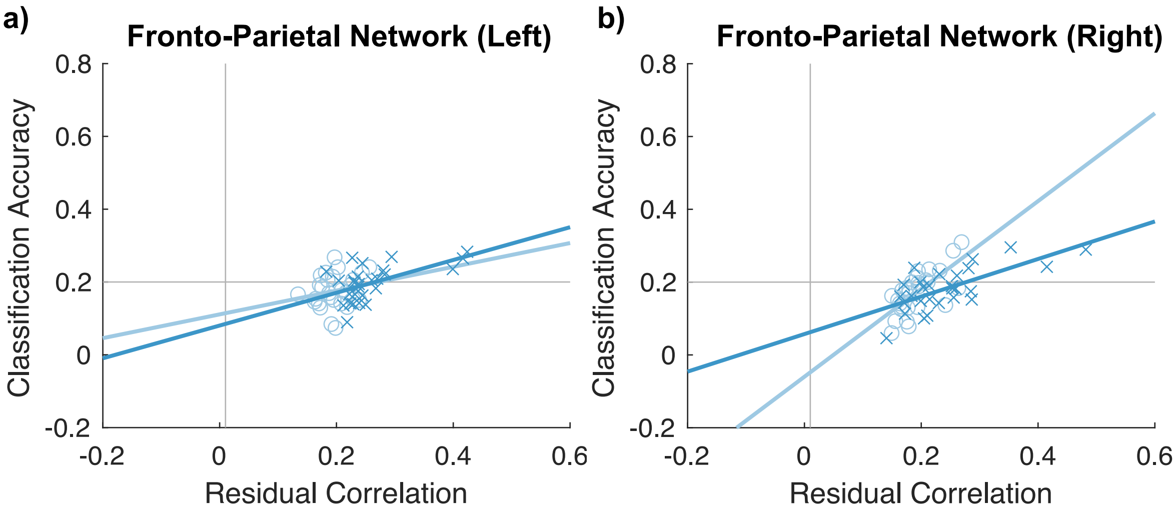


c)

| Hemisphere | N | Variable | Estimate | Std. Error | t value | *p* |
|---|---|---|---|---|---|---|
| Left | 28 | Intercept | 0.065 | 0.073 | 0.888 | 0.379 |
| | | Residual Correlations | 1.282 | 0.370 | 3.467 | **0.001*** |
| | | Condition | 0.055 | 0.045 | 1.220 | 0.228 |
| | | Residual Correlations x Condition | -0.580 | 0.237 | -2.466 | **0.018*** |
| Right | 28 | Intercept | 0.024 | 0.062 | 0.385 | 0.702 |
| | | Residual Correlations | 1.448 | 0.326 | 4.437 | **4.768 x $10^{-5}$*** ** |
| | | Condition | 0.051 | 0.038 | 1.329 | 0.190 |
| | | Residual Correlations x Condition | -0.527 | 0.205 | -2.578 | **0.013*** |

**Supplementary Figure 9. Linear regression relating classification accuracy and residual correlations in Glasser Atlas ROIs overlapping the fronto-parietal network (FPN) in the left (A) and right (B) hemispheres.** Each point is one Glasser ROI colored by condition (dark-colored X's: attended; light colored O's: ignored). Table in (C) depicts the linear regressions relating classification accuracy and residual correlations in attended and ignored conditions in the fronto-parietal network. *p<.05, **p<.01, ***p<.001. Bold values represent those passing correction for multiple comparisons (2 models).

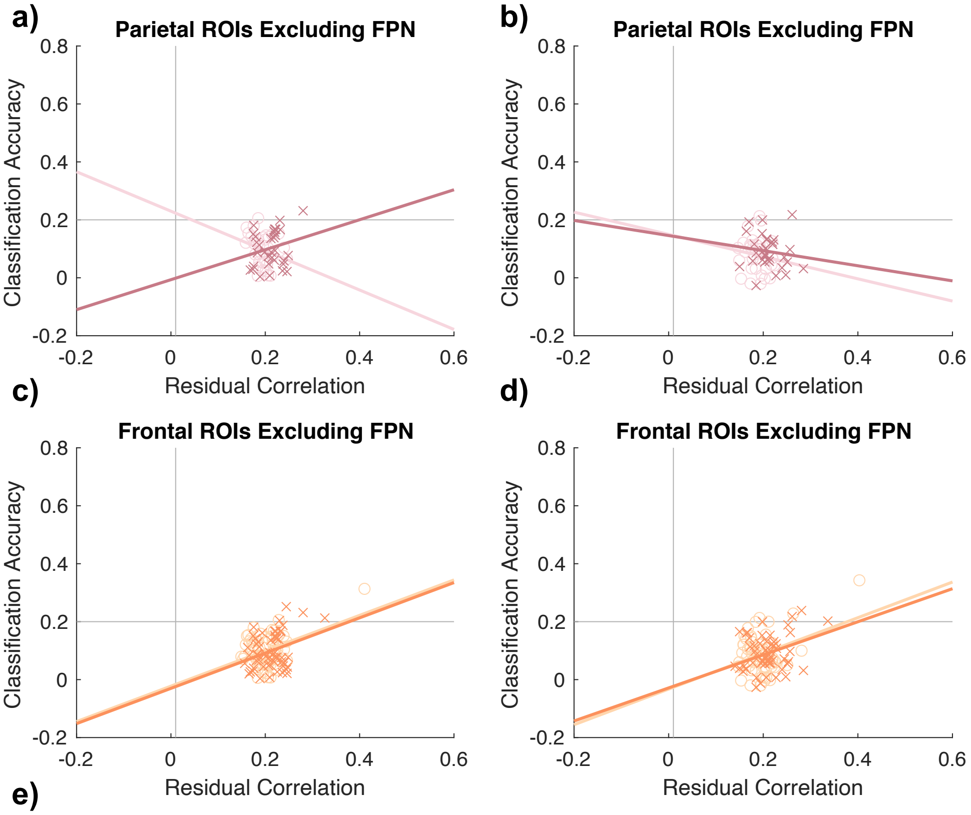


e)

| Lobe | Hemisphere | N | Variable | Estimate | Std. Error | t value | *p* |
|---|---|---|---|---|---|---|---|
| Parietal | Left | 34 | Intercept | 0.199 | 0.018 | 11.342 | **$5.858 \times 10^{-17}$***** |
| | | | Residual Correlations | 0.281 | 0.150 | 1.872 | 0.066 |
| | | | Condition | 0.003 | 0.011 | 0.258 | 0.798 |
| | | | Residual Correlations x Condition | -0.188 | 0.099 | -1.901 | 0.062 |
| | Right | 34 | Intercept | 0.244 | 0.018 | 13.245 | **$5.318 \times 10^{-20}$***** |
| | | | Residual Correlations | -0.094 | 0.177 | -0.530 | 0.598 |
| | | | Condition | -0.026 | 0.011 | -2.332 | 0.023* |
| | | | Residual Correlations x Condition | 0.022 | 0.110 | 0.202 | 0.840 |
| Frontal | Left | 59 | Intercept | 0.146 | 0.025 | 5.898 | **$3.833 \times 10^{-8}$***** |
| | | | Residual Correlations | 0.878 | 0.203 | 4.317 | **$3.390 \times 10^{-5}$***** |
| | | | Condition | 0.015 | 0.016 | 0.885 | 0.378 |
| | | | Residual Correlations x Condition | -0.306 | 0.139 | -2.204 | 0.030* |
| | Right | 59 | Intercept | 0.152 | 0.023 | 6.485 | **$2.371 \times 10^{-9}$***** |
| | | | Residual Correlations | 0.882 | 0.203 | 4.337 | **$3.130 \times 10^{-5}$***** |
| | | | Condition | 0.013 | 0.015 | 0.892 | 0.374 |
| | | | Residual Correlations x Condition | -0.339 | 0.130 | -2.618 | **0.010*** |

**Supplementary Figure 10. Linear regression relating classification accuracy and residual correlations in Glasser Atlas ROIs from the Parietal (A and B) and Frontal (C and D) lobes, excluding Glaser ROIs that overlap the fronto-parietal network (FPN).** Data are depicted separately for the left (A and C) and right (B and D) hemispheres. Each point is one Glasser ROI colored by condition (dark-colored X's: attended; light colored O's: ignored). Table (E) depicts the linear regressions relating classification accuracy and residual correlations in attended and ignored conditions. *p<.05, **p<.01, ***p<.001. Bold values represent those passing correction for multiple comparisons (4 models).
